# Epigenetic memory of temperature sensed during somatic embryo maturation in 2-year-old maritime pine trees

**DOI:** 10.1101/2024.06.27.600784

**Authors:** J.-F. Trontin, M.D. Sow, A. Delaunay, I. Modesto, C. Teyssier, I. Reymond, F. Canlet, N. Boizot, C. Le Metté, A. Gibert, C. Chaparro, C. Daviaud, J. Tost, C. Miguel, M.-A. Lelu-Walter, S. Maury

**Affiliations:** BioForBois, FCBA, Pôle Industrie Bois & Construction, Cestas, 33610, France; P2e, Université d’Orléans, INRAE, EA 1207 USC 1328, 45067 Orléans, France; Biosystems and Integrative Sciences Institute, Faculdade de Ciências, Universidade de Lisboa, 1749-016 Lisboa, Portugal; BioForA, INRAE, ONF, UMR 0588, 45075 Orléans, France; Sylviculture Avancée, FCBA, Pôle Ressources Forestières des Territoires, Cestas, 33610, France; IHPE, Université de Perpignan, UMR 5244, 66100, Perpignan, France; Laboratory for Epigenetics and Environment, Centre National de Recherche en Génomique Humaine, CEA - Institut de Biologie François Jacob, Université Paris Saclay, 91000 Evry, France

**Keywords:** *Pinus pinaster*, somatic embryogenesis, memory, epigenetics, development, heat/cold stress, DNA methylation, methylome, sequence capture bisulfite

## Abstract

Embryogenesis is a brief but potentially critical phase in the tree life cycle for adaptive phenotypic plasticity. Using somatic embryogenesis in maritime pine, we found that temperature during the maturation phase affects embryo development and post-embryonic tree growth for up to three years. We examined whether this somatic stress memory could stem from temperature- and/or development-induced changes in DNA methylation. To do this, we developed a 200 Mb custom sequence capture bisulfite analysis of genes and promoters to identify differentially methylated cytosines (DMCs) between temperature treatments (18, 23, and 28°C) and developmental stages (immature and cotyledonary embryos, shoot apical meristem of 2-year-old plants) and investigate if these differences can be mitotically transmitted from embryonic to post-embryonic development (epigenetic memory). We revealed a high prevalence of temperature-induced DMCs in genes (8-14%) compared to promoters (less than 1%) in all 3 cytosine contexts. Developmental DMCs showed a comparable pattern but only in the CG context, and with a high trend towards hypo-methylation, particularly in the promoters. A high percentage of DMCs induced by developmental transitions were found memorized in genes (up to 45-50%) and promoters (up to 90%). In contrast, temperature-induced memory was lower and confined to genes after both embryonic (up to 14%) and post-embryonic development (up to 8%). Using stringent criteria, we identified ten genes involved in defense responses and adaptation, embryo development and chromatin regulation that are candidates for the establishment of a persistent epigenetic memory of temperature sensed during embryo maturation in maritime pine.

**Graphical abstract:** 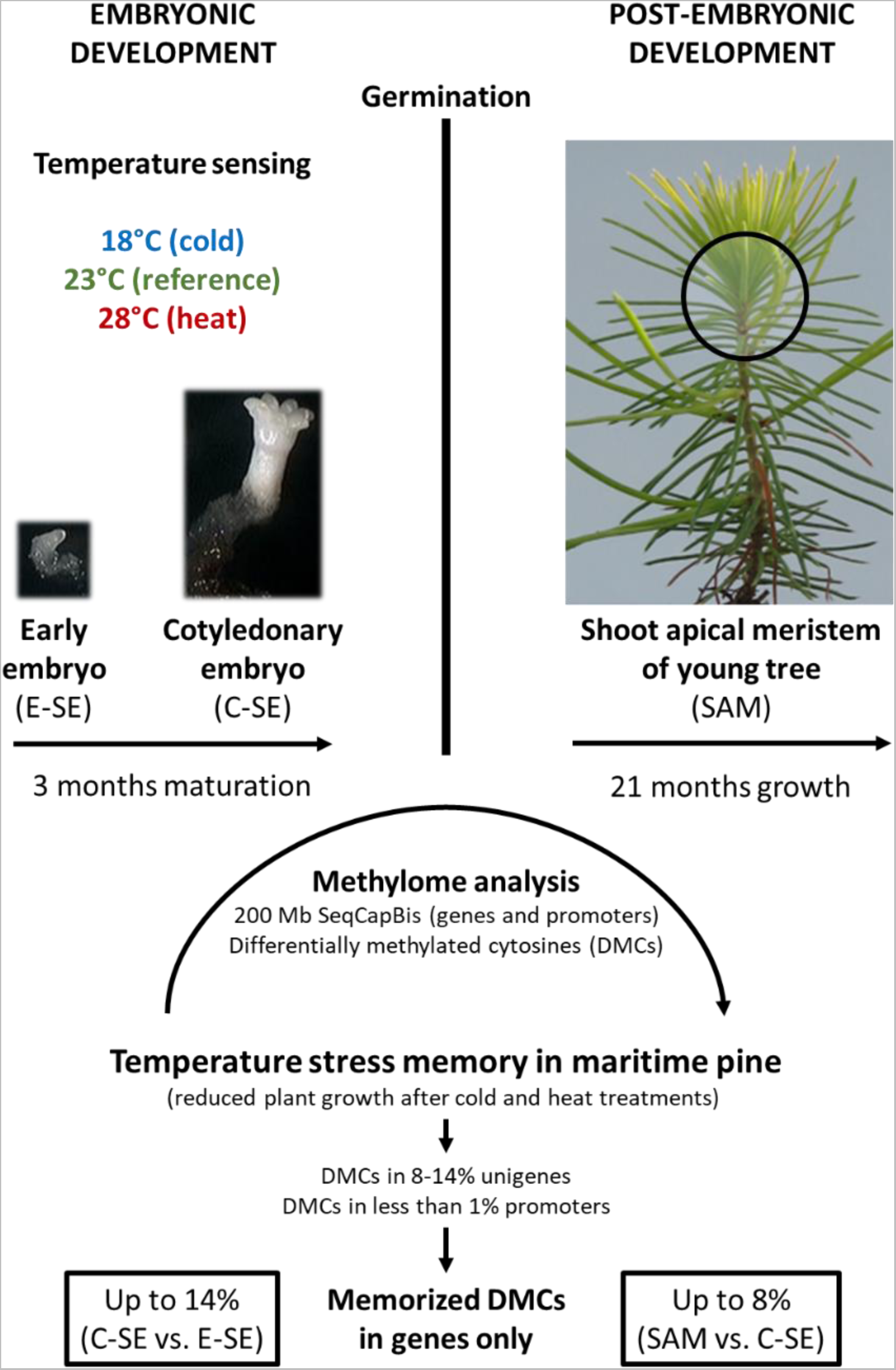

## INTRODUCTION

Due to their longevity, large size, and late reproductive phase, adaptability of trees to climate change is a major concern. Projections all point towards a significant rise in temperature (Adak et al. 2023) that will expose most forests to recurrent and/or more severe heat and drought episodes (Plomion et al. 2016, Hammond et al. 2022). Such environmental stresses can have adverse effects on tree capacity to produce seeds (Clark et al. 2021). In maritime pine (*Pinus pinaster* Ait.) for example, a major plantation conifer in the Mediterranean basin, a strong decline in seed production has been observed in French orchards since the late 2000s (Boivin and Davi 2016). Biotic and abiotic factors are suspected, including temperature effects on flowering and seed formation.

Embryogenesis is a short phase of the tree life cycle resulting in embryo formation within seed. It occurs from the fertilized egg cell (zygotic embryo, ZE), and more rarely from unfertilized reproductive or differentiated somatic cells (somatic embryo, SE). Somatic embryogenesis is a promising vegetative propagation way for conifers (Klimaszewska et al. 2016). In these species, embryogenesis involves a cascade of auxin- and abscisic acid (ABA)-mediated events, from pro-embryogenesis (ZE) to early and late embryogenesis (ZE, SE), coordinating apical-basal and radial patterning (Trontin et al. 2016a, von Arnold et al. 2016). Auxin-mediated cell fate decisions result in early delineation of primary shoot (SAM) and root (RAM) apical meristems followed by procambium at the early cotyledonary embryo stage (Palovaara et al. 2010, Brunoni et al. 2019).

There is increasing evidence that post-embryonic meristems may have a fundamental role in plant adaptation and memory as primary sensors of environmental stresses (Lämke and Bäurle 2017, Maury et al. 2019, Zhu et al. 2023). The same could apply to embryonic meristems and any embryogenic cell (Castander-Olarieta et al. 2021, Trontin et al. 2021).

Besides genetics, plant adaptation may also operate through either adaptive phenotypic plasticity (i.e., the ability of a genotype to express different phenotypes), or robustness (when a genotype shows a rather stable phenotype). These processes could support rapid evolutionary changes and acclimation of plants (Nicotra et al. 2010) and are thought to involve epigenetic factors affecting the expression of genes, but not their nucleotide sequence (Maeji and Nishimura 2018, Zhu et al. 2023). Such changes can be reversibly imprinted in the genome at a much higher rate than genetic mutations (Sow et al. 2018) through development (e.g., bud break, Conde et al. 2017; dormancy, Kumar et al. 2016; embryogenesis, Markulin et al. 2021; aging, Li et al. 2023) and environmental effects (e.g., drought, Jacques et al. 2021; heat, Perella et al. 2022). They are mostly transient modifications allowing the resetting of expression patterns at key developmental stages and continuous adaptation to new conditions (Hemenway and Gehring 2023). Part of these changes can be stably maintained by cell division and support the neoformation of adapted organs (Maury et al. 2019). Epigenetic changes can even promote genetic variation leading to local adaptation (Sáez-Laguna et al. 2014, Platt et al. 2015, Alakärppä et al. 2018). As SAM alternately produce vegetative and sexual organs, any stable (epi)genetic modification of meristematic cells could be transmitted to gametes and progenies (Hofmeister et al. 2020).

Somatic stress memory has been reported in annuals (Jacques et al. 2021, Zhu et al. 2023) and perennials (de Freitas Guedes et al. 2018, Tan 2023). Current evidence points to synergistic control by epigenetic and transcription factors (Liu et al. 2021, Gao et al. 2022b) that could be critically expressed in meristems (Birnbaum and Roudier 2017, Maury et al. 2019), especially in the case of thermomorphogenesis (Zhu et al. 2023). A memory of temperature during embryogenesis with long-lasting effects was reported in Norway spruce (Johnsen et al. 2005, Kvaalen and Johnsen 2008, Skrøppa 2022). Genetic selection has been ruled out (Besnard et al. 2008) and evidence for an epigenetic control emerged from transcriptomics (Yakovlev et al. 2011, 2014) and profiling of small RNAs (Yakovlev et al. 2010, 2020, Yakovlev and Fossdal 2017).

DNA methylation has a pivotal role in epigenetics and has been the focus of numerous studies to explore inheritable phenotypic variation (Seymour and Becker 2017). It predominantly occurs in plants at cytosine sites as 5-methylcytosine (5mC) in the CG, CHG (H: A, C, or T) and CHH contexts (Stroud et al. 2014, Zhang et al. 2018). Different DNA methyltransferase classes are involved in methylation maintenance at these sites (MET1, CMT3, and CMT2, respectively) while *de novo* methylation is mediated by DRM1/2 through RNA-directed DNA methylation (RdDM) involving 24-nucleotide short interfering RNAs (siRNAs, Matzke and Mosher 2014). There are both functional similarities and divergence of these pathways in higher plants (Ausin et al. 2016, Niu et al. 2022, Li et al. 2023) that may explain different DNA methylation patterns in some groups (e.g., higher levels in all 3 contexts for gymnosperms).

Cytosine (de)methylation can occur at high rate in the genome (Yao et al. 2021) and has long been associated in angiosperms with biological processes, cellular functions and regulation of gene expression during embryonic (Ji et al. 2019, Chen et al. 2020, Wójcikowska et al. 2020, Markulin et al. 2021) and post-embryonic development (Zhang et al. 2018), symbiotic interactions (Vigneaud et al. 2023), stress responses and somatic memory (Zhang et al. 2018, Le Gac et al. 2018, Liu and He 2020, Rajpal et al. 2022). Similar essential roles are anticipated for gymnosperms with also a focus on genome stability as these species typically show much larger and heavily methylated genomes because of high content in transposable elements (Ausin et al. 2016, Niu et al. 2022).

In maritime pine, transcriptomics suggested that epigenetic reprogramming is occurring during embryogenesis (de Vega-Bartol et al. 2013, Rodrigues et al. 2018, 2019). There is evidence for both methylation maintenance during early embryogenesis and *de novo* RdDM towards the late stages. Global DNA methylation changes or methylation-sensitive amplification polymorphisms have been detected during somatic embryogenesis in conifers (Miguel et al. 2016) and associated with embryo developmental stages (Teyssier et al. 2014) or maturation ability (Klimaszewska et al. 2009). Global changes were also observed in pine following temperature sensing during SE initiation or maturation suggesting that DNA methylation could contribute to the establishment of an epigenetic stress memory (Castander-Olarieta et al. 2020, Pereira et al. 2021). Despite increasing availability of genomic resources in trees (Plomion et al. 2016, Sterck et al. 2022), evidence for temperature effect on genes involved in DNA methylation (Yakovlev et al. 2016), and the possibility for bisulfite sequencing, it is still unknown what is the extent of 5mC imprinting during conifer embryogenesis as a result of both developmental transition and temperature sensing effects.

In this work, we used somatic embryogenesis as an *in vitro* process mimicking zygotic embryogenesis in maritime pine (Morel et al. 2014a, Trontin et al. 2016b, Rodrigues et al. 2019) to investigate whether developmental transitions and temperature during SE maturation could induce changes in phenotype and methylome. We produced early (E-SE) and cotyledonary (C-SE) embryos at 18, 23 (reference) and 28°C and further regenerated somatic plants (Fig. 1A). We used available pine genomic resources to perform a targeted bisulfite sequencing of genes and promoters (Fig. 1B) in E-SE, C-SE (direct development or temperature effects) and the SAM of young trees (remaining, delayed effects). We found large sets of genes but fewer promoters containing differentially methylated cytosines (DMCs) induced by embryonic (comparing C-SE and E-SE) and post-embryonic (SAM vs. C-SE) development or temperature in E-SE, C-SE and SAM in response to cold (18 vs. 23°C) or heat (28 vs. 23°C). A significant part of development-(in genes and promoters) but also temperature-induced DMCs (in genes only) were mitotically transmitted from the embryonic to the post-embryonic phase. We could demonstrate both developmental and stress epigenetic memory established during embryogenesis in maritime pine.

**Figure 1:**
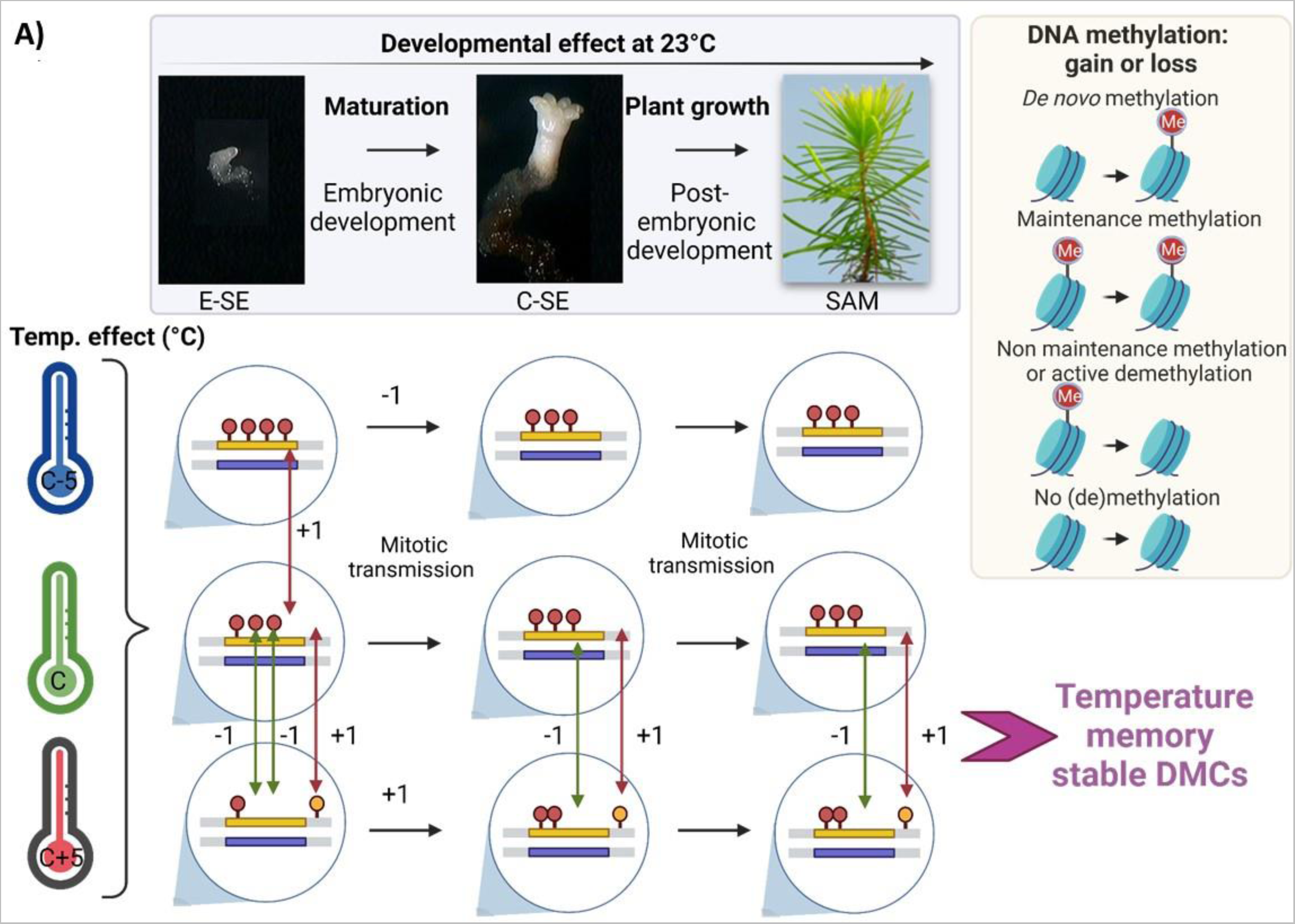
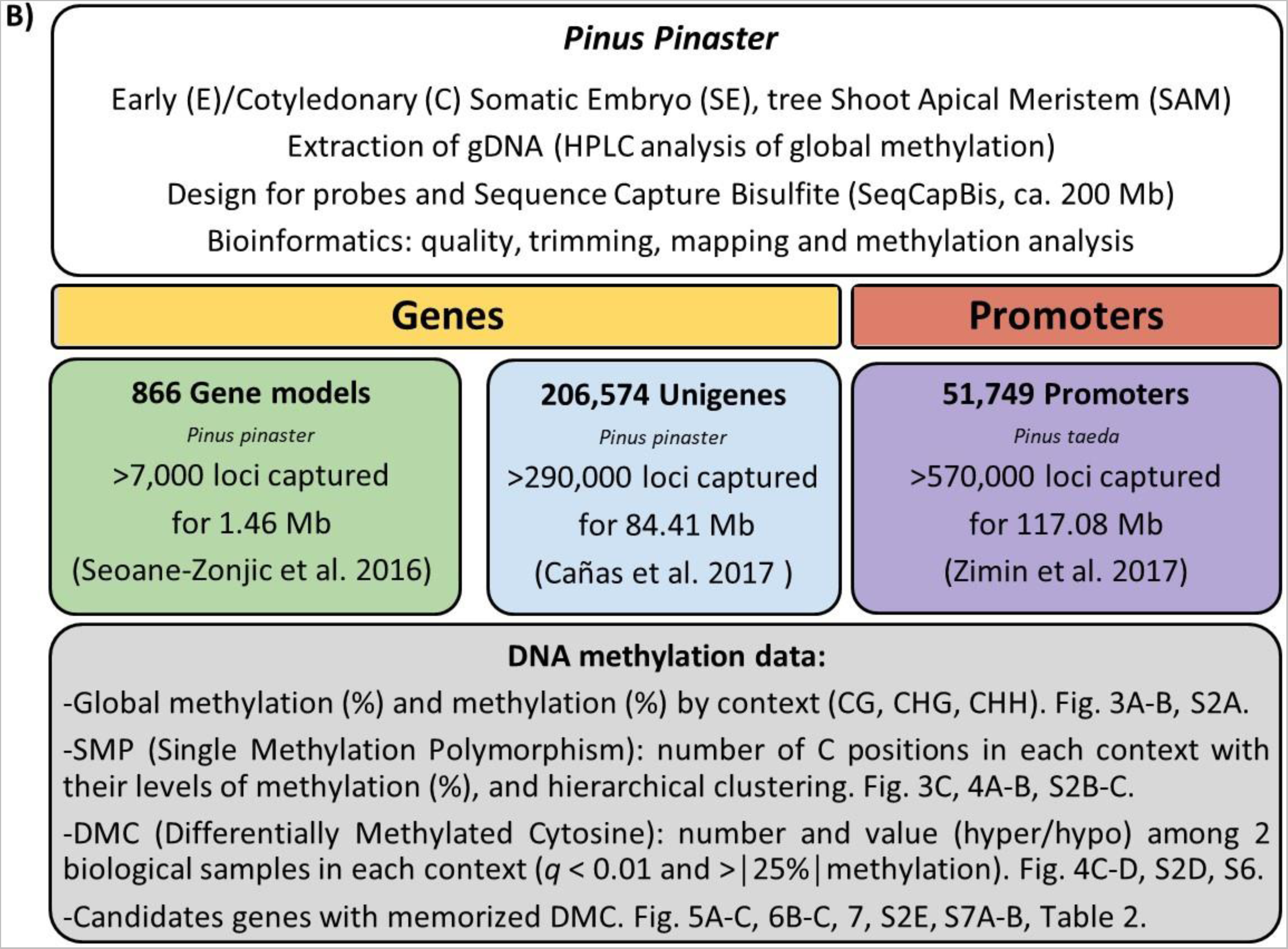
Experimental designs for production of biological material (**A**) and DNA methylation analyses (**B**). **A.** The PN519 embryogenic line was reactivated from the cryopreserved stock (2 weeks) and propagated until the embryogenic masses were fully established (8 additional weeks), i.e., consisting of multiple, actively multiplying immature somatic embryos (SE). This initial step of the embryonic phase of development (early embryogenesis) was performed at the reference control temperature (C = 23°C). Embryogenic masses were then subjected to 3 temperature treatments (18, 23 and 28°C, i.e., C-5, C, and C+5) during the whole maturation phase to produce early SE (E-SE, after 1 week) and cotyledonary SE samples (C-SE, after 11-17 more weeks). After storage at 4°C for 4-5 months (embryo post-maturation treatment and synchronization), the C-SE were germinated at 23°C (3 weeks) and the resulting plantlets were gradually acclimated (4 months; 23-25°C) and further grown in greenhouse conditions (16 months; 15-30°C) to obtain collectable shoot apical meristems (SAM). At this stage of the post-embryonic development, somatic plants were 21 months old since germination (juvenile vegetative phase). E-SE, C-SE and plants were phenotyped at various times around the sampling date for methylome analysis to study temperature effects (cold: 18 vs. 23°C; heat: 28 vs. 23°C) as well as development effects (at 23°C). This fully clonal design allowed testing for temperature- and/or development-induced memory from embryonic (E-SE vs. C-SE, heterotrophic state) to post-embryonic phase (C-SE vs. SAM, photosynthetically competent state). For this purpose, DNA methylation analyses were performed at cytosine sites to identify gain (+1) or loss (-1) in 5 methylcytosine (5mC). Such methylation marks were checked for mitotic transmission between developmental stages. **B.** Genomic DNA (gDNA) was extracted from E-SE, C-SE, and SAM samples for both HPLC (global DNA methylation) and sequence capture bisulfite (methylome) analyses. After sequence quality control using bioinformatic tools, over 867,000 loci were successfully captured in gene models (> 7,000 loci), unigenes (> 290,000 loci) and promoters (> 570,000 loci). We analyzed the global DNA methylation level, the percentage of methylation by context, the occurrence of single methylation polymorphisms (SMPs) and differentially methylated cytosines (DMCs). We finally identified candidate genes with memorized DMCs at the embryonic or post-embryonic phase.

## MATERIAL AND METHODS

### Plant material, experimental design, and sampling

We investigated one cryopreserved *P. pinaste*r embryogenic line (PN519) initiated in 1999 from an immature seed (G0.4304*G0.4301; pedigree: Landes Forest, France). It is a gold genotype to study embryo development (Lelu-Walter et al. 2016, Trontin et al. 2016b, Llebrés et al. 2018).

PN519 was subjected to three temperature treatments (18, 23, 28°C; Fig. 1A) during the maturation step enabling the development of immature E-SE into C-SE. 23°C is the reference temperature used during somatic embryogenesis in maritime pine (Trontin et al. 2016b). To study the effect on DNA methylation at lower (18°C) or higher (28°C) maturation temperature than the reference (23°C), we sampled and characterized three types of biological materials: i) E-SE after 1 week maturation; ii) C-SE after at least 12 weeks maturation; and iii) SAM collected from 21-month-old plants. The production process of this plant material is presented in Fig. 1A.

For each type of embryonic (E-SE, C-SE) or post-embryonic (SAM) material and temperature treatment combination, 3-5 biological replicates were made, immediately frozen in liquid nitrogen and stored at -80°C until processing. E-SE sample (400 mg fresh mass, f.m.) consisted of multiple immature embryos attached to remaining embryogenic tissue. C-SE sample (200 mg f.m.) included 231-306 single embryos separated from the residual tissue. SAM sample (2-4 mg f.m.) resulted from the dissection of the apical meristem of a single shoot apex.

### Culture of somatic embryos and plants

Reactivation from the cryopreserved stock, multiplication, and maturation of PN519 were performed in Petri dishes (94x16 mm) containing 23.5 mL semi-solid mLV basal medium (Litvay et al. 1985, Klimaszewska et al. 2001) and closed by cling film (2 rounds). Cultures were incubated at a selected temperature (± 1°C) in darkness (TC175S, Aqualytic, Dortmund, Germany).

Embryogenic tissues were thawed at 37°C, drained and plated on filter paper (Whatman N°2, 70 mm), then placed on mLV supplemented with 0.5 M sucrose, 2 µM 2,4-dichlorophenoxyacetic acid (Sigma-Aldrich/Merck KGaA, Darmstadt, Germany), 1 μM 6-benzyladenine (Duchefa Biochemie, Haarlem, The Netherlands) and solidified with 4.5 g/L gellan gum (HP696, Kalys, Bernin, France). After 24 h, the filters supporting cells were transferred to the same reactivation medium, but containing 0.3 M sucrose. After 48 h, the cells were scraped off the filter and transferred to the same medium with standard sucrose (0.09 M) and gellan gum concentrations (3 g/L Gelrite, Duchefa Biochemie) for multiplication. From 2 weeks after thawing, the reactivated line was weekly subcultured on multiplication medium to promote rapid growth. When propagating easily, embryogenic tissue was suspended in liquid maturation medium (mLV supplemented with 0.2 M sucrose, 80 µM ABA, Ecochem, China), and distributed on filter paper (Whatman N°2, 70 mm) at a cell density of 50-300 mg f.m./filter. The filter was then placed onto maturation medium solidified with high gellan gum (9 g/L Gelrite) and subcultured once on fresh medium after 4 weeks. At 23°C, C-SE development typically occurs after 10-14 weeks (Morel et al. 2014a).

C-SE were collected under the binocular and stored in darkness at 4°C on a modified mDCR medium (Gupta and Durzan 1985) without plant growth regulator, 0.175 M sucrose and 9 g/L gellan gum (Gelrite, Duchefa Biochemie). After 4-5 months, C-SEs were germinated (N = 120/condition) on mDCR containing 58 mM sucrose and 4.6 g/L gellan gum (4 g/L Gelrite, Duchefa Biochemie; 0.6 g/L HP696, Kalys). After 3 weeks at 23°C, a subset of viable embryos (N = 96/condition) were transferred in 8-ml miniplugs (15% peat, 85% coco fiber, Preforma plug trays, ViVi, Mijlweg, The Netherlands) and cultivated in a growth room (1 month, 23°C), then in the greenhouse (3 months, 25°C) until full acclimatization. Young trees were transplanted in horticultural substrate at age 4 (110 ml container) and 15 months (one-l pot) and further grown in the greenhouse until age 25 months. At this time, a subset of trees (N = 45/condition) were transferred to the nursery until planting in fall at age 31 months (Oct. 2018). The field trial consists of 3 blocks with a fully randomized design (N = 5 trees/condition within each block).

### Phenotypic characterization

Embryogenic cultures matured at the 3 temperatures were characterized (Mat. S1) through macro- and micro-morphological qualitative observations (behavior of cultures, overall time required to produce C-SE) and quantitative measurements (embryo yield, mass, size, and morphology).

Embryo yield was calculated as the number of C-SE harvested per gram f.m. embryogenic tissue matured. It was estimated after 12-18 weeks maturation depending on temperature treatment (see Results). Mean C-SE f.m. (mg) was estimated from the 5 samples collected for DNA methylation analysis by dividing the total mass of each sample (ca. 200 mg) by the number of embryos harvested. Embryo size (mm) and morphology were investigated based on pictures (N = 60/condition) analyzed with the Acrobat Reader DC (Adobe) measurement tool. We assessed total embryo length (from the hypocotyl base to the tip of the largest cotyledon), cotyledon ring length (from the insertion point on the hypocotyl to the tip of the largest cotyledon), hypocotyl length (from the root pole base to the insertion point of the cotyledon ring) and width (just below insertion point of the cotyledon ring), and the number of cotyledons.

We also investigated the delayed effect of maturation temperature on C-SE development, from germination (viability and germination rates) to plant survival and growth (height, height increase, terminal bud elongation). Embryo viability and germination rates were checked after 3 weeks, i.e., just before transfer to miniplugs. Viability rate was calculated as the percentage of viable C-SEs with elongated hypocotyl and cotyledons on germination medium. Germination rate is the percentage of viable C-SEs with root development (checked under the binocular).

Plant survival in the greenhouse conditions was recorded at ages 5, 8, and 15 months after germination and then yearly in the field (2019, 2020, 2021). Plant height (cm) was measured at ages 8, 15, 36, and 65 months. Relative height increase (%) was calculated as (H_n_-H_n-1_/H_n-1_)*100 with H_n_ the total height observed at measurement n. To characterize bud “break” during early spring, which is an indefinite, temperature-dependent process in maritime pine (from bud swelling to elongating bud and prickly shoot), we monitored the terminal bud length (from main plant axis) in late March and early May during the spring of 2019 and 2021. Relative increase in length (%) was calculated as (L_n_-L_n-1_/L_n-1_)*100 with L_n_ the length at measurement n.

### Quantification of soluble carbohydrates and starch

E-SE and C-SE samples (n = 5 biological replicates) were lyophilized and ground into fine powder using a MM400 Retsch mixer Mill. Each sample (4-20 mg dry mass, d.m.) was extracted three times at 85°C in 1 mL ethanol:water (80:20, v/v) for soluble carbohydrates and starch following Bonhomme et al. (2010) modified by Gautier et al. (2019). Mannitol was added in the extracts as an internal standard (0.25 mg/mL). Pooled, purified, and dried supernatants were suspended in 250 µL ultrapure water and centrifuged before analyses by HPLC (see Gautier et al. 2019). Soluble carbohydrates were identified by co-elution with standards and quantified from the calibration curves (mg carbohydrates/g d.m, Mat. S1). From the resulting pellets, starch content was quantified in glucose equivalents (mg glucose/g d.m., Mat. S1) after hydrolysis with amyloglucosidase (Morel et al. 2014a). Each sample was assayed one (soluble carbohydrates) or two times (starch).

### Total protein assay

Total protein extracts were prepared in five replicates for each sample type from frozen material (25-50 mg f.m.) according to Morel et al. (2014a). Protein concentration (µg/mg f.m., Mat. S1) was determined using the Bradford assay (1 time) with bovine serum albumin as a standard.

### DNA extraction and global DNA methylation percentages by HPLC

Genomic DNA (gDNA) was extracted from E-SE (400 mg f.m.), C-SE (200 mg f.m.) and individual SAM (n = 3 for each developmental stage and temperature condition; 27 samples overall) using a CTAB protocol (Doyle and Doyle 1987) and was stored at -80°C. gDNA quantity and quality were assessed using a NanoDrop spectrometer (Thermo Fisher Scientific, Waltham, MA, USA). For estimating global DNA methylation, gDNA was enzymatically hydrolyzed into nucleosides and analyzed by HPLC (Zhu et al. 2013, Genitoni et al. 2020).

### Sequence capture bisulfite for methylome analysis

Designed probes are available in Mat. S2. An equimolar pool of 1 μg gDNA extracted from 3 biological samples was made for each developmental stage in each temperature treatment. Bisulfite treatment and capture using our custom designed probes were made on these 9 types of biological samples according to SeqCap Epi Target Enrichment System (SeqCap Epi Developer XL Enrichment Kit) using Roche recommendations (https://sequencing.roche.com/content/dam/rochesequence/worldwide/resources/brochure-seqcap-epi-SEQ100146.pdf). Similar equimolar pools of 0.4 μg gDNA at ca. 20 ng/µL were used for sequencing at the CNRGH (Evry, France) with paired ends (2×150 bp) on an Illumina HiSeq4000 platform following. Raw data were stored in FASTQ files with a minimal theoretical coverage of 100X.

A Roche NimbleGen SeqCap EZ Design (https://sequencing.roche.com/content/dam/rochesequence/worldwide/resources/brochure-seqcap-ez-prime-choice-probes-SEQ100193.pdf) of custom 200 Mb was performed by NimbleGen service using the 3 following supporting reference sequences and conditions (see Fig. 1B and Mat. S2 for details): i) 866 *P. pinaster* gene models from Seoane-Zonjic et al. (2016) with the condition of covering the whole gene body sequences, ii) 206,574 *P. pinaster* unigenes derived from Cañas et al. (2017) with the conditions of covering the whole unigene (gene body) sequences if·size is under 1,578 bp, and 789 bp in 5’ and 3’ for a total of 1,578 bp if unigene size is over or equal to 1,578 bp, and iii) 51,749 promoters from Pita v2.0 *Pinus taeda* genome (https://treegenesdb.org/FTP/Genomes/Pita/v2.0/; Zimin et al. 2017) with the condition of covering until 1,205 bp upstream to the transcription starting site (TSS).

The bioinformatic pipeline for methylome analysis was adapted from the ENCODE pipeline (https://www.encodeproject.org/wgbs/) and installed on the Galaxy instance, accessible at IHPE (http://galaxy.univ-perp.fr/, Perpignan, France) according to Sow et al. (2021) and Dugé de Bernonville et al. (2022). Single Methylation Polymorphism (SMP) data were processed against the 3 sets of supporting reference sequences (Fig. 1B) using the methylKit R package (Akalin et al. 2012) to identify Differentially Methylated Cytosines (DMCs) with a minimum coverage of 10X, differential methylation between two samples of at least 25% for all contexts and a *q*-value < 0.01. Gene annotation was achieved by similarity search using megaBLAST alignments (BLAST+ BLASTN v2.10.1) against *Arabidopsis thaliana* genome v11 available at TAIR database (https://www.arabidopsis.org/). Gene Ontology (GO) enrichment analysis of methylated genes was performed using default parameters (see Mat. S3) of Metascape (Zhou et al. 2019, https://metascape.org/gp/index.html#/main/step1).

### Statistical analyses of phenotypic data

Biological data were analyzed using Statview 5.0 (SAS Institute Inc). Differences in means (reported with 95% confidence limits) between temperature treatments for quantitative traits were assessed by analysis of variance (ANOVA). Student-Newman-Keul’s (SNK) post-hoc test was used to identify differences between groups (*p* < 0.05) when ANOVA indicated significant effects. Biochemical data were similarly assessed by one-way ANOVA and multiple comparison of means with Tukey contrasts (*p* < 0.05) using the R version 3.3.2 (R Development Core Team 2011, R: A Language and Environment for Statistical Computing. Vienna: R Foundation for Statistical Computing). Statistical tests and *p-*values are indicated as recommended by Wasserstein and Lazar (2016).

## RESULTS

### The maturation temperature affects biological and biochemical aspects of embryonic development and post-embryonic growth

The first effect observed after only 1 week maturation of the PN519 line was the temperature-dependent increase in cell proliferation (Fig. 1A, Fig. 2A). For high temperatures such as 28°C, cell proliferation could be modulated by reducing inoculum density from 100 (standard) to 50 mg/filter (Fig. 2A). At these 2 plating densities, both lower (18°C) and higher temperature (28°C) than the reference (23°C) significantly delayed embryo development (Table S1).

**Figure 2:**
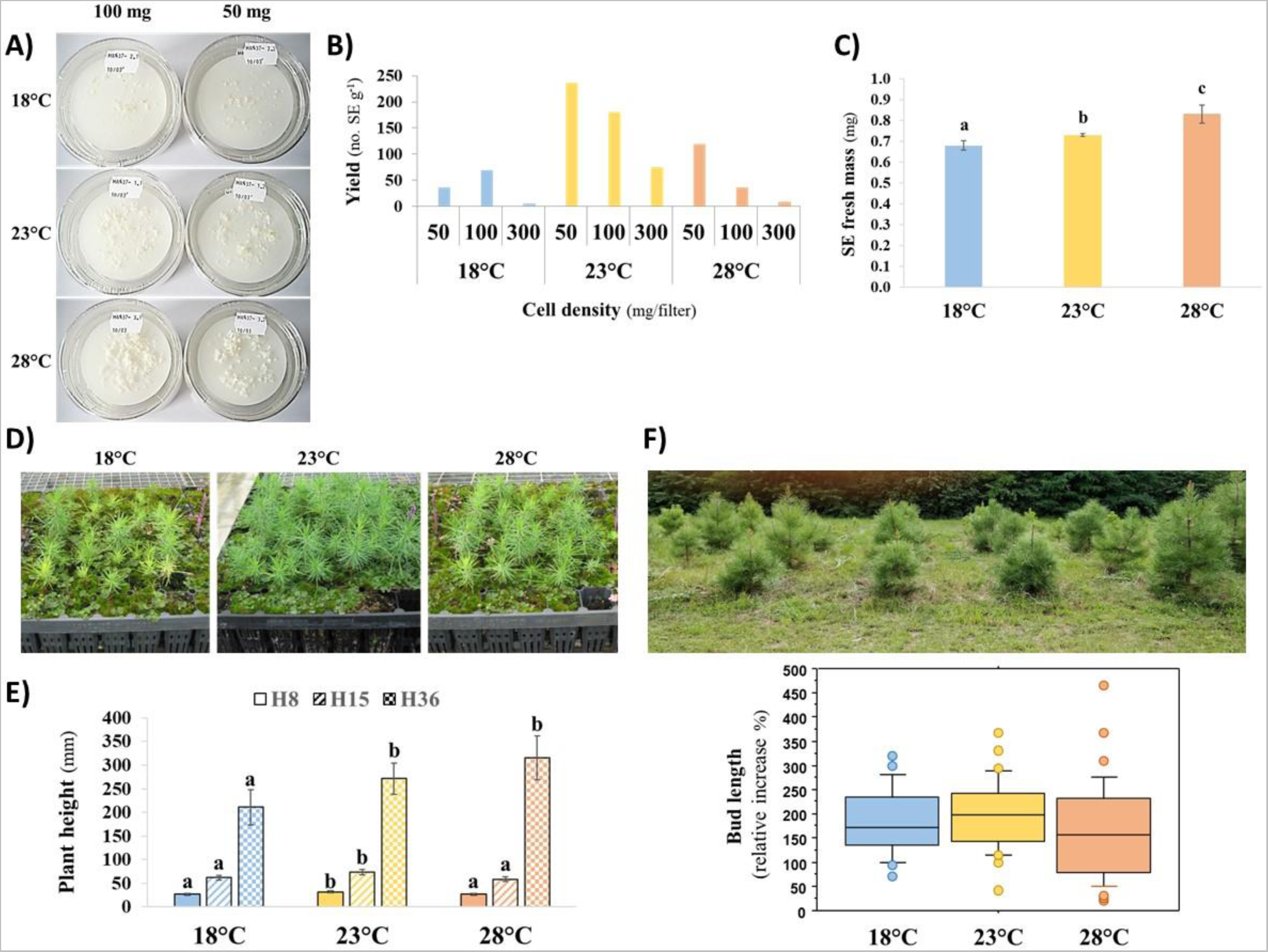
Effect of temperature during maturation (18, 23, 28°C) on embryonic and post embryonic development of PN519 somatic embryos. **A.** Observed embryogenic tissue proliferation after 1 week maturation at 2 cell densities (100 and 50 mg / filter). Note that cell growth increases with temperature. **B.** Yield in cotyledonary somatic embryos (C-SE) according to initial cell density (50-300 mg/filter) and temperature after 12-13 (23°C), 15-16 (23°C) or 17-18 weeks (18°C) maturation time. **C.** Mean fresh mass of cotyledonary somatic embryos (C-SE). Bars: 95% confidence limits. Significant differences (*p* < 0.05) are indicated by different letters. The difference observed between 18 or 23°C and 28°C is significant at *p* < 0.01. **D.** Somatic plant behavior 15 months after germination. Note the apparent, general yellowing (chlorosis) of the plant batches from embryos treated at 18°C and 28°C compared to the standard, greener batch (23°C). **E.** Mean plant height at age 8 (H8), 15 (H15) and 36 months (H36). Growth in the greenhouse (H8, H15) or at field (H36). Bars: 95% confidence limits. For each variable, significant differences are indicated by different letters at *p* < 0.01 (H8, H15) or 0.05 (H36). **F.** Relative increase in length of plant terminal bud during spring 2019 (age 37-38 months) and 2021 (age 61-62 months). Box plot displaying the 10th, 25th, 50th, 75th, and 90th percentiles and outliers. ANOVA did not detect any significant effect between year and temperature conditions. Picture: a view of the field trial at age 65 months.

E-SE samples were produced at a high cell density in the 3 temperature conditions. In contrast, C-SE samples were obtained at standard cell density at 18°C and 23°C and reduced cell density at 28°C to avoid tissue overgrowth. Yields in C-SE were computed after 12-13 (23°C), 15-16 (28°C) or 17-18 weeks (18°C) maturation at the 3 selected cell densities to produce E-SE and C-SE samples. They were the highest at 23°C (Fig. 2B). Despite downward adjustment of cell density and/or extension of maturation time, embryo yields were lower at 28°C, and even more at 18°C.

The mean fresh mass of individual C-SE harvested for DNA methylation analysis significantly increased (*p* < 0.05) with temperature, from 0.68 (18°C) to 0.83 mg (28°C) (Fig. 2C). The increase is greater between 28 and 23°C (*p* < 0.01) than between 23 and 18°C (*p* < 0.05).

Significant effects of temperature on C-SE size (Fig. S1A) were detected for total length (*p* = 0.0001), hypocotyl length, and width (*p* < 0.0001). Since ANOVA did not detect any effect for cotyledon ring length, it can be deduced that it is mainly the hypocotyl that accounts for the observed differences. Post-hoc SNK tests confirmed that hypocotyl size was significantly reduced (*p* < 0.01) at 18°C compared to 23/28°C for both length (0.79 vs. 1.16 / 1.09 mm) and width (0.59 vs. 0.67 / 0.70 mm). ANOVA also revealed a temperature effect on cotyledon number (p = 0.0008). Embryos bore fewer cotyledons (*p* < 0.01) at 18°C (3.0) than at 23°C (3.7) or 28°C (3.9).

Quantitative biochemical differences were detected between E-SE and C-SE (Fig. S1B), i.e., more proteins (Table 1, Mat. S1), sucrose, starch, and oligosaccharides of the raffinose family (RFOs, raffinose, stachyose) in C-SE, but less glucose and fructose (Fig. S1B). In addition, maturation temperature affected the content of starch and soluble carbohydrates in both E-SE and C-SE (Fig. S1B). In E-SE, starch was higher at 18°C (47.4 mg/g d.m.) compared to 23 and 28°C (19.7-28.8 mg/g) and glucose increased as a function of temperature, from 131.4 mg/g (18°C) to 199.0 mg/g (28°C). In C-SE, starch accumulated more efficiently at 23°C (127.2 mg/g) than at 18°C (103.9 mg/g) or 28°C (86.0 mg/g), and sucrose content was lower at 28°C (99.5 vs. 121.3 mg/g).

**Table 1:** Main phenotypic and methylome characteristics of somatic embryos (embryonic phase) and corresponding plants (post-embryonic phase) after different temperature treatments (18, 23 and 28°C) during maturation (conversion of early embryos to cotyledonary embryos). Temperature effects are presented separately from the development effects studied at the reference temperature (23°C, see Fig. 1A for experimental design). C-SE: Cotyledonary Somatic Embryos; DMC: Differentially Methylated Cytosine; E-SE: Early Somatic Embryos; GO: Gene Ontology; RFOs: Raffinose Family Oligosaccharides; SAM: plant Shoot Apical Meristem; SE: Somatic Embryo; SMP: Single Methylation Polymorphism.

| Study scale | Temperature effects (18, 23 or 28°C)<br>(cold: 18 vs. 23°C; heat: 28 vs. 23°C) | Development effects (23°C)<br>(embryonic: E-SE vs. C-SE; post-embryonic: C-SE vs. SAM) |
| --- | --- | --- |
| **Phenotype**<br>(embryo growth, development, quality)<br>Fig. 2, S1<br>Tables S1, S2 | E-SE proliferation increases with temperature. Fig. 2A.<br>Cold and heat delayed C-SE development. Table S1.<br>Temperature affects C-SE yield (lower at 18/28°C), fresh mass (increasing from 18 to 28°C), size (smaller hypocotyl length/width at 18°C), and morphology (fewer cotyledons at 18°C). Fig. 2B, 2C, S1A.<br>No significant differences in total protein content (Mat. S1).<br>Higher starch content in E-SE at 18°C and in C-SE at 23°C. Fig. S1B.<br>Glucose concentration increased in E-SE with temperature. Sucrose content is lower at 28°C in C-SE. Fig. S1B.<br>Disturbed plant physiology during post-embryonic growth after treatment at 18°C (intense yellowing) and 28°C (moderate). Fig. 2D.<br>Lower plant height at 8-15 (18/28°C) and 36 months (18°C) originating in the early stages of post-embryonic growth. Fig. 2E, S1D.<br>No effect on C-SE germination and viability, plant survival (age 5 to 15 months), height (65 months), and terminal bud elongation in the field (37-62 months). Fig. S1C, S1E, 2F, Table S2. | More proteins in C-SE than E-SE (39 vs. 4 µg/mg f.m., Mat. S1)<br>C-SE accumulates more starch than E-SE. Fig. S1B<br>C-SE contains more sucrose and RFOs (raffinose, stachyose) but less glucose and fructose than E-SE. Fig. S1B. |

**Table 1.** Continued
| Study scale | Temperature effects (18, 23 or 28°C)<br>(cold: 18 vs. 23°C; heat: 28 vs. 23°C) | Development effects (23°C)<br>(embryonic: E-SE vs. C-SE; post-embryonic: C-SE vs. SAM) |
| --- | --- | --- |
| **Global methylome**<br>(levels and SMPs)<br>Fig. 3, S2-5 | No significant differences in global DNA methylation ($p = 0.179$) related to development. Fig. S2A.<br>Hierarchical clustering based on SMP occurrence does not discriminate properly between temperatures (18, 23, 28°C). Fig. S2C, S3B. | Significant differences in global DNA methylation related to development (E-SE < C-SE, $p = 0.014$). Fig. S2A.<br>SMP-based clustering of development stages (E-SE, C-SE, SAM) in CHH > CHG > CG and promoter > gene orders. Fig. S2C, S3B. |
| **Differential methylome**<br>(DMCs)<br>Fig. 3, 4, S3, S7-9 | Genes: SMP prevalence in significant temperature- or remaining temperature-DMCs is higher in CG and CHG (3-11%) than CHH (1-5%) contexts. Fig. 3C, S3D.<br>Promoters: SMP prevalence in significant temperature- or remaining temperature-DMCs is considerably lower than in genes (< 0.1 % in all contexts). Fig. 3D.<br>Temperature-DMCs have similar hyper-/hypo-methylation ratios in all contexts. They are most variable for the CHG context (0.308-7.652) compared to CG (0.474-2.321) and CHH (0.333-1.577), and promoters (0.308-7.652) compared to genes (0.286-1.751). Fig. S7.<br>12% of unigenes (ca. 25,000 out of 206,574) but less than 0.4% of promoters (< 200 out of 51,749) contain temperature- or remaining temperature-DMCs in each context for E-SE, C-SE and SAM. Fig. 4A.<br>1,500-3,000 (CG) and 1,700-9,500 (CHG/CHH) unigenes contain at least 5 temperature-DMCs. The highest values (8,000-9,000) are observed for C-SE under cold or heat (CHH). Strikingly, no promoters contain at least 5 temperature-DMCs. Fig. S8A.<br>GO enrichment analysis of unigenes/promoters containing DMCs ($\geq 5$) highlights functions in primary/secondary metabolisms, response to heat and temperature stimulus, especially in promoters. Fig. S9. | Genes: SMP prevalence in significant development-DMCs is higher in CG and CHG (6-11%) than CHH (2-5%) contexts. Fig. 3C, S3D.<br>Promoters: compared to genes, SMP prevalence in significant development-DMCs is lower in CG (< 0.1%) and CHG (< 1.3%) but similar in CHH (1.3-3.1%) contexts. Fig. 3D.<br>Development-DMCs are mostly hypomethylated in genes (ratios: 0.082-0.929) and promoters (0.003-0.446), notably in CHH/CHG contexts (0.003-0.041). Hypomethylation affects both embryonic (0.014-0.929) and post-embryonic DMCs (0.003-0.724). Fig. S7.<br>12% of unigenes (ca. 25,000) contain development-DMCs and 8-14% promoters in the CHH context (4,000-7,000), but less than 4% in CHG (< 2,000) and 0.4% (< 200) in CG. Fig. 4B.<br>2,500-3,000 (CG) to 7,000-8,500 (CHG/CHH) unigenes contain at least 5 development-DMCs with similar figures at the embryonic and post-embryonic phases. This is far fewer promoters and only in CHG (50-60) and CHH (619-1605) contexts. Fig. S8B.<br>GO enrichment analysis of unigenes/promoters containing DMCs ($\geq 5$) reveals functions in primary/secondary metabolisms and also, intriguingly, response to temperature stimulus. Fig. S9. |

**Table 1. End**
| Study scale | Temperature effects (18, 23 or 28°C)<br>(cold: 18 vs. 23°C; heat: 28 vs. 23°C) | Development effects (23°C)<br>(embryonic: E-SE vs. C-SE; post-embryonic: C-SE vs. SAM) |
| --- | --- | --- |
| **Memory methylome**<br>(conserved DMCs)<br>Fig. 4-6, S3<br>Table 2 | • Temperature-DMCs can be transmitted from embryonic (E-SE to C-SE) to post-embryonic (C-SE to SAM) phases. Fig. 4C, Fig. 5.<br>• Genes: levels of temperature-induced memory (% memorized DMCs) during the embryonic phase are higher in CG/CHG (10-14%) than CHH (3-5%) context. Post-embryonic levels are lower in CG/CHG (2-8%) but similar in CHH context (1-5%). Fig. 5B, S3E.<br>• Promoters: no temperature-induced memory could be detected after both embryonic and post-embryonic development. Fig. 5B.<br>• There are more genes containing memorized DMCs for cold (4000-8000) than for heat (2000-5000) after both embryonic and post-embryonic development (Fig. 4C).<br>• GO enrichment analysis of temperature-induced memory in unigenes unravel molecular functions related with stress response (mainly heat, UV, jasmonic acid, beta-alanine metabolism) and carbohydrate metabolisms (glycolysis/gluconeogenesis, starch and sucrose). Fig. 6.<br>• Annotated genes containing most memorized DMCs ($\geq 5$) at the post-embryonic phase (CSE to SAM) relate to proteins involved in defense/adaptation responses (cold/heat: Cystein-rich receptor-like; cold: DAD1-like lipase, Cytochrome P450; heat: Heat shock protein, ABC transporter), chromatin regulation (heat: Histone H3.1), and embryo development (cold: Subtilisin-like; heat: Embryo defective MAPK kinase, arabinogalactan methylesterase, insulinase). Table 2. | • Development-DMCs (identified in C-SE vs. E-SE) can be transmitted to the post-embryonic phase (C-SE to SAM). Fig. 5.<br>• Genes: levels of development-induced memory (% memorized DMCs) observed after the post-embryonic phase (C-SE to plant SAM at 23°C) reached 45-50% in CG/CHG contexts and 25-42% in CHH context. Fig. 5B, S3E.<br>• Promoters: compared to genes, levels of development-induced memory are similar in CHH (18%) or higher for CG (90%) and CHG (61%) contexts. Fig. 5B. |

The C-SE viability and germination rates were estimated after 4-5 months storage (Fig. S1C). Despite the observed morphological and biochemical differences (Fig. 2C, S1A, S1B), C-SE showed similar viability and germination rates when entering the post-embryonic phase (3 weeks germination), whatever the maturation temperature. Viability rates were high and in a narrow range 85-89%. Germination rates were lower and displayed a wider range (50-58%), but differences were not significant. Similarly, we did not observe any major effect of maturation temperature on plant survival after 5-, 8-, and 15-months (Table S2). A slightly lower survival rate was observed after 15 months at 23°C (73%) compared to 18°C (81%) or 28°C (80%). However, the distribution of dead plants within the seedling trays was uneven and points to undetected plant management issues. Furthermore, we noticed at age 15 some substantial chlorosis of plants obtained from embryos matured at 18°C (intense) and 28°C (moderate) as compared to 23°C (Fig. 2D). Accordingly, these two plant batches showed significantly lower height at ages 8 and 15 months (Fig. 2E). This difference originated from early stages of post-embryonic growth (before 8 months) since the three plant lots showed no significant variation in relative height increase between 8 and 15 months (Fig. S1D), a period that essentially matches spring shooting. Significant differences in growth were still detected after field planting (Fig. 2E). Three years after germination, the 18°C plant batch showed a lower average height. No significant difference could be detected after about 3 years of field growth (age 65 months, Fig. S1E). However, the trial suffered from significant game damages (35% of plants) during the first year. Similarly, we did not detect delayed effects when considering terminal bud elongation during early spring of 2019 and 2021 (age 37-62 months, Fig. 2F).

In summary, lower (18°C) or higher (28°C) temperature than the reference (23°C) during embryo maturation adversely affects embryonic development and post-embryonic plant growth up to 36 months following germination.

### Whole genome DNA methylation is more impacted by developmental transition than temperature during embryo maturation

To study DNA methylation in our biological material (E-SE, C-SE, SAM) following embryo maturation at 18, 23 or 28°C (9 types of samples), we performed a global DNA methylation analysis by HPLC and a custom methylome profiling using SeqCapBis (Fig. 1B).

Global DNA methylation values ranged from 15.1% (E-SE, 23°C) to 29.3% (C-SE, 23°C) (Fig. S2A) with a significant developmental effect (ANOVA, *p* = 0.013). DNA methylation levels were found lower in E-SE compared to C-SE (*p* = 0.014) with intermediate, not significantly different values for SAM samples. No temperature effect was observed (ANOVA, *p* = 0.179).

A capture probe design for a 200 Mb SeqCapBis of the *P. pinaster* methylome including genes models (866), unigenes (206,574) and promoters (51,749) was used for DNA methylation analysis (Fig. 1B, Mat. S2). Mapping coverage (Table S3) ranged from a mean of 19x (promoters) to 57x (unigenes), and 235x (gene models). Considering unigenes and promoters, cytosine methylation levels ranged from 65.0-71.6% (CG) to 55.6-58.5% (CHG), and 10.4-20.7% (CHH) (Fig. S2B, Fig. S4). Unigenes display higher CG (*p* = 0.035) and lower CHH methylation levels (*p* = 0.009) than promoters. Gene models and unigenes followed similar methylation patterns in the 3 contexts (Fig. S3A, Fig. S4).

The number of cytosine positions that could be analyzed for Single Methylation Polymorphisms (SMPs) is ascending from CG (0.13-0.99 million) to CHG (0.21-1.72) and CHH (0.87-5.89) contexts, as expected due to their genomic frequency (Fig. 3A,B, Fig. S3C). In line with the capture design (Fig. 1B), we found more SMPs in unigenes than promoters (6-8 times, Fig. 3A,B) or gene models (30-40 times, Fig. 3A and Fig. S3C). Using SMP values, correlations between samples (Fig. S5) were found higher for promoters (*r* up to 0.93 in CG, 0.88 in CHG, 0.78 in CHH) than for unigenes (*r* up to 0.88 in CHH, 0.86 in CG, 0.80 in CHG). Hierarchical clustering of samples based on SMP occurrence (Fig. S2C and S3B) was quite consistent with developmental stages in CHH (promoters, unigenes), CHG (unigenes, promoters) and CG (promoters) contexts. E-SE generally cluster with SAM rather than C-SE samples suggesting deep methylation rearrangements during development. In contrast, there was no influence of the temperature treatments on the clustering.

**Figure 3:**
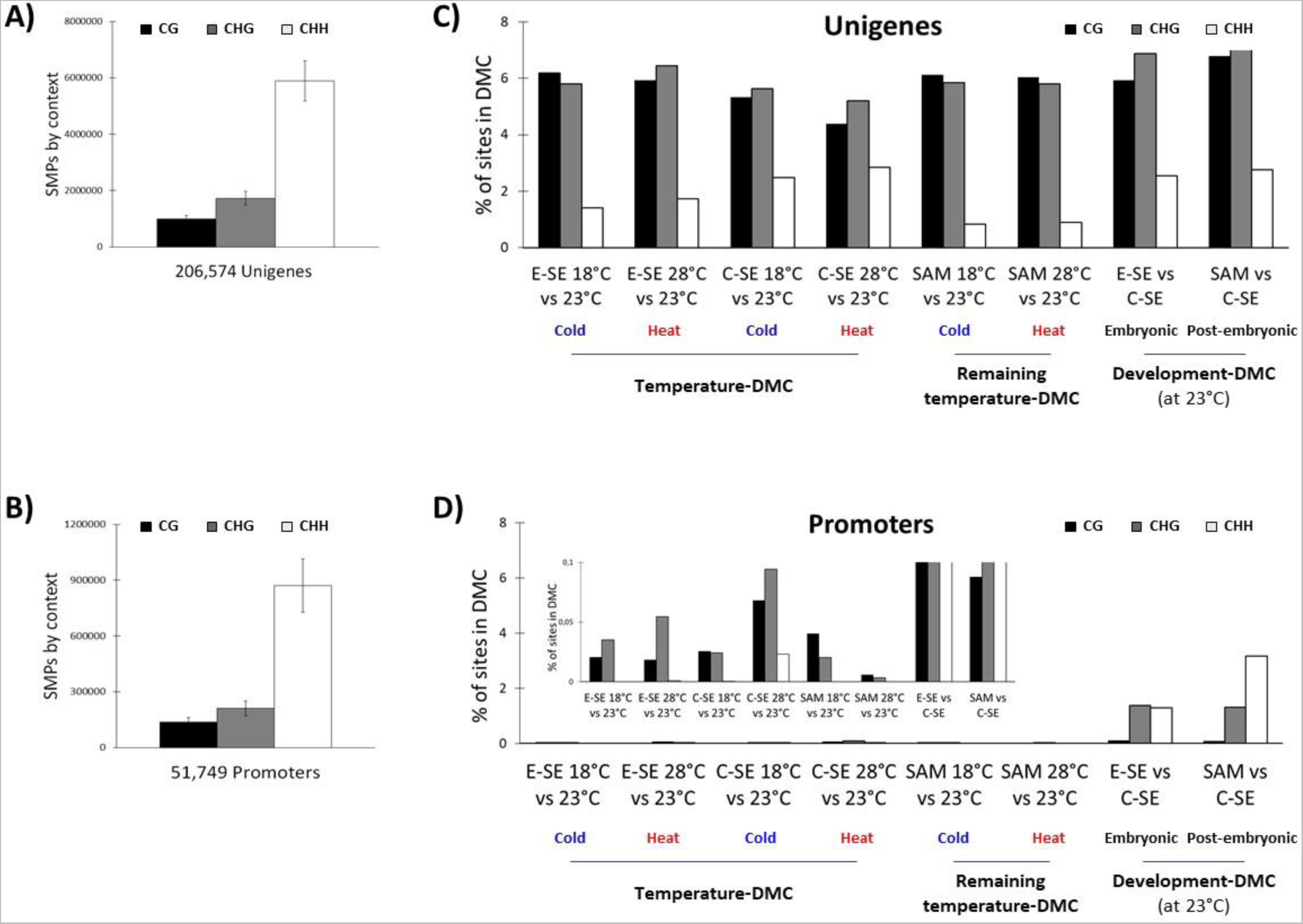
Differential Methylation. **A-B.** Number of Single Methylation Polymorphisms (SMPs) detected in the 3 contexts of methylation (CG, CHG, CHH). **A**. Unigenes; **B**. Promoters. **C-D**. Percentage of SMP sites in significant Differentially Methylated Cytosines (DMCs), i.e., with *q*-value < 0.01 and at least a 25% methylation difference, compared to the total number of SMP sites analyzed by cytosine context (CG, CHG, CHH) for all *P. pinaster* samples. **C**. Unigenes; **D**. Promoters. The different types of DMC (cold/heat temperature-, remaining temperature- and embryonic/post-embryonic development-DMC) are indicated (see Fig. S6).

### High prevalence of differentially methylated cytosines in genes compared to promoters, especially in response to temperature

Differentially Methylated Cytosines (DMCs) were identified in the 3 methylation contexts and for the 3 designs (gene models, unigenes, and promoters) among temperature treatments for cold (18 vs. 23°C) and heat effect (28 vs. 23°C), or among developmental stages at 23°C for direct (E-SE vs. C-SE) and delayed effect (C-SE vs. SAM). DMCs were grouped into three classes (Fig. S6): i) temperature-DMCs identified in E-SE or C-SE exposed to maturation temperature (i.e., during embryonic development), ii) remaining temperature-DMCs identified in plant SAM (i.e., after post-embryonic development) and iii) development-DMC identified after embryonic (C-SE) or post-embryonic development (plant SAM). For genes (unigenes and gene models), 3-11% of CG/CHG and only 1-5% of CHH SMP sites were in significant temperature- and remaining temperature-DMCs in all conditions (Fig. 3C, Fig. S3D). A similar pattern was observed for development-DMCs (6-11% of CG/CHG, and 2-5% of CHH SMP sites). In contrast, for promoters (Fig. 3D), less than 0.1% of SMPs in all 3 contexts were in temperature- and remaining temperature-DMCs while development-DMCs were also lower than in genes for CG (< 0.1%) and CHG (< 1.3%), but similar in CHH (1.3-3.1%).

Temperature- and remaining temperature-DMCs in unigenes, gene models, and promoters were similarly hyper- and hypo-methylated in all 3 contexts (Fig. S7). In contrast, both embryonic and post-embryonic development-DMCs were found mostly hypo-methylated in genes and even more in promoters, notably in CHH/CHG contexts (Fig. S7, Table 1).

About 25,000 out of the 206,574 unigenes (12%) had at least one temperature-, remaining temperature- or development-DMC in each context (Fig. 4A,B). Filtering for more than 5 DMCs resulted in less than 3,000 (1.5%) unigenes in the CG context and 8,000-10,000 in the CHG/CHH context (4-5%) considering both temperature (Fig. S8A, highest values for C-SE under heat or cold in CHH) and development effects (Fig. S8B, similar numbers for embryonic and post-embryonic development). GO enrichment analysis (see Mat. S3) showed that the main molecular functions (-log10(P)>6) of unigenes containing temperature-, remaining temperature- or development-DMCs are related to primary (photosynthesis, polysaccharide metabolism, glycolysis/gluconeogenesis, nucleobase-containing small molecule metabolic process) or secondary metabolisms (phenylpropanoid metabolic process) and protein autophosphorylation (see Fig. S9B for heat effect on C-SE, cold remaining effect on SAM and post-embryonic developmental effect). Response to heat and other abiotic stresses (reactive oxygen species, osmotic, etc.) were also well represented (-log10(P)>4), especially for temperature, but also development effects. In both cases molecular functions associated with embryo and seedling development could be identified.

**Figure 4:**
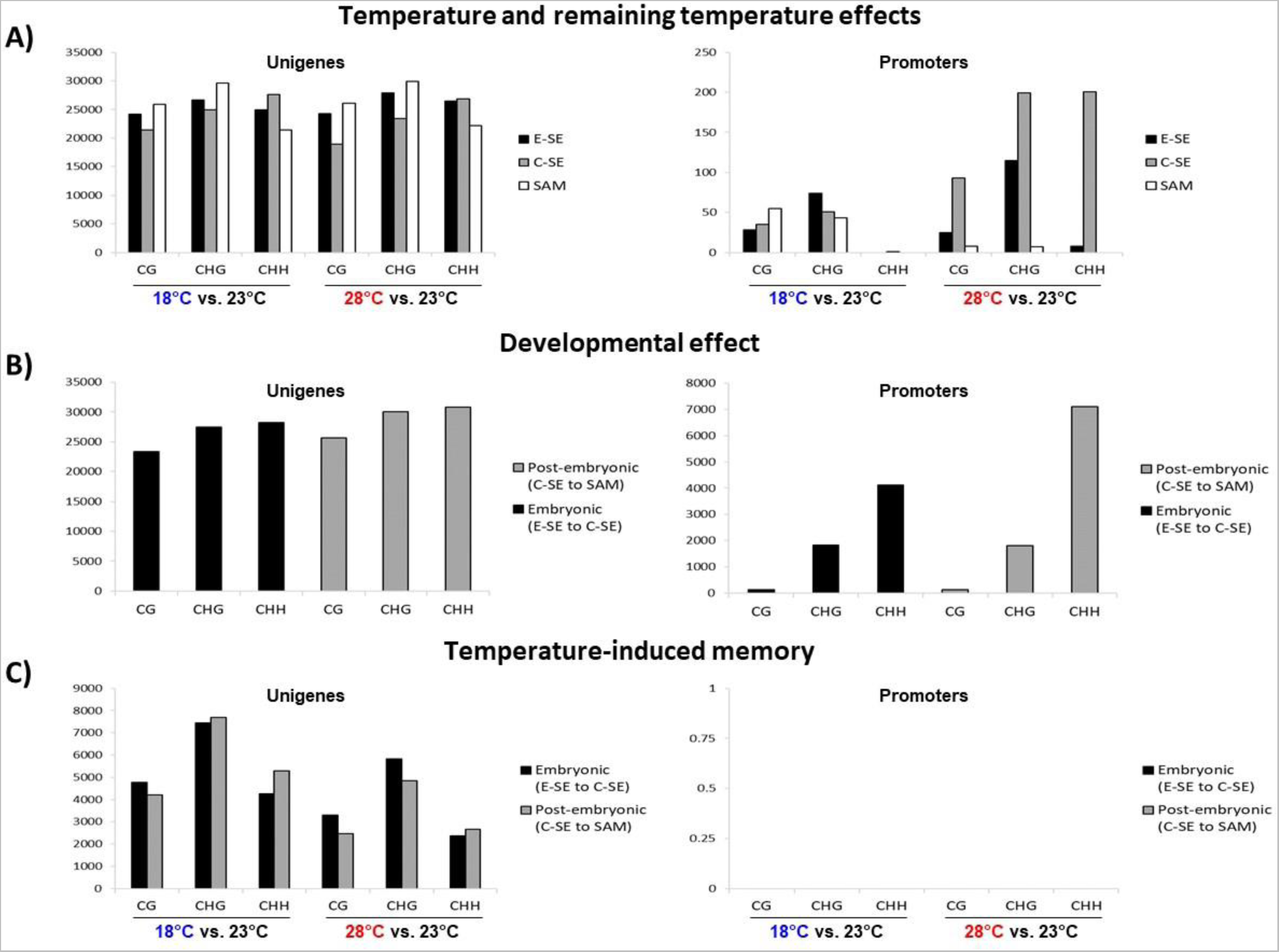
Number of unigenes (left) and promoters (right) with Differentially Methylated Cytosines (DMCs) in the 3 contexts (CG, CHG, CHH). **A.** Temperature effect (seen in E-SE, C-SE) and remaining temperature effect (seen in SAM) following cold (18 vs. 23°C) and heat (28 vs. 23°C) treatments. **B.** Developmental effect (at 23°C) during embryonic (E-SE to C-SE) and post-embryonic development (C-SE to SAM). **C.** Temperature-induced memory during embryonic (E-SE to C-SE) and post-embryonic development (C-SE to SAM) following cold (18 vs. 23°C) and heat (28 vs. 23°C) treatments.

Regarding promoters, less than 200 out of the 51,749 promoters (0.4%) displayed temperature- or remaining temperature-DMCs in each context, with highest values for C-SE under heat (CHG/CHH) and the lowest for SAM, especially under heat in the 3 contexts (Fig. 4A). In contrast, development-DMC were found in up to 2,000-7,000 promoters (10 to 35 times more, 4-14%), notably in the CHH context, with the highest number for post-embryonic development (Fig. 4B). As previously observed for unigenes, GO enrichment analysis (Mat. S3) confirmed for promoters molecular functions related to primary and secondary metabolic processes as well as specific “response to temperature stimulus” for C-SE under heat (- log10(P)>4) and, intriguingly, also post-embryonic developmental effects (-log10(P)>6, Fig. S9A). In contrast, limited evidence for molecular functions associated with embryo or seedling development was found in promoters.

### DMCs can be stably transmitted from the embryonic to the post-embryonic phase with higher occurrence when induced by developmental transitions compared to temperatures

In order to test if DMCs induced by temperature (cold or heat) or embryo development (at 23°C) during SE maturation can be mitotically transmitted to support an epigenetic memory, each DMC class (temperature-, remaining temperature- and development-DMC) was compared to a similar DMC class obtained at another developmental stage (i.e., E-SE vs. C-SE, C-SE vs. SAM for temperature-DMCs, embryonic vs. post-embryonic phase for development-DMCs). Identical DMCs (i.e., found at the same position) with the same hyper-/hypo-methylation status were then identified (Fig. 5A). In the case of unigenes for example, heat (28 vs. 23°C)-induced CG DMCs in E-SE (54,079) were compared to heat-induced CG DMCs in C-SE (40,068) and the 7,232 identical DMCs were filtered for similar methylation status (both hyper- or hypo- methylation). In this way, we identified 4,749 stable, memorized temperature-DMCs (heat) during embryo maturation (11.9% of memorized DMCs, Fig. 5B). Similarly, we found 46,718 identical DMCs comparing CG DMCs induced by embryonic development at 23°C (C-SE vs. E-SE) and CG DMCs induced by post-embryonic development (SAM vs. C-SE). After filtration, we identified 23,155 memorized development-DMCs (50% of memorized DMCs).

**Figure 5:**
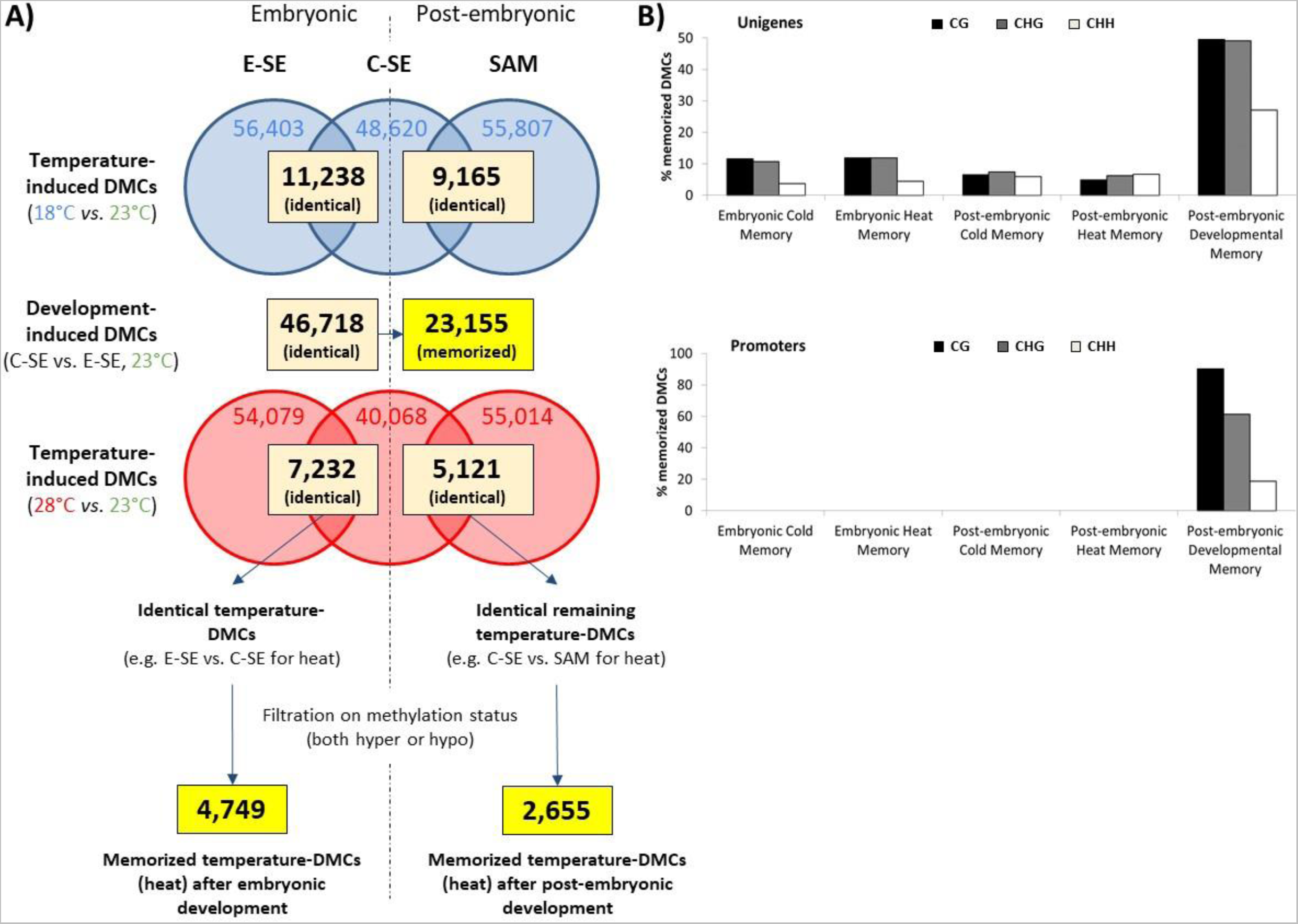
Memory methylome. **A.** Analysis flow chart to estimate the number of stable, memorized DMCs (highlighted in yellow) induced by temperature (cold: 18 vs. 23°C; heat: 28 vs. 23°C) or embryo development (C-SE vs. E-SE at 23°C) during maturation for unigenes in CG context. We first calculated the number of identical DMCs, i.e., that are found at the same positions between different sample types: i) Temperature effect: number of identical DMCs induced by cold or heat comparing E-SE and C-SE (embryonic, temperature-DMCs) or C-SE and SAM (post-embryonic, remaining temperature-DMCs). ii) Development effect: number of identical DMCs induced by embryo development comparing C-SE (embryonic development-DMCs) and SAM (post-embryonic development-DMCs). Each category of identical temperature- or development-DMCs were then filtered for the same differential of hyper- or hypo-methylation to identify stable, memorized DMCs. We found e.g., 4,749 or 2,655 memorized heat-induced DMCs after embryonic or post-embryonic development, respectively, and 23,155 development-induced DMCs memorized after post-embryonic growth. **B.** Percentage (%) of memorized DMCs (all 3 contexts CG, CHG, CHH) in unigenes (up) or promoters (down) induced by temperature (cold: 18 vs. 23°C; heat: 28 vs. 23°C) or embryo development (C-SE vs. E-SE at 23°C). Post-embryonic developmental memory and both embryonic and post-embryonic temperature-induced (cold or heat) memories are shown.

Overall, the percentage of memorized temperature-DMCs (cold or heat) in unigenes (Fig. 5B) and gene models (Fig. S3E) was higher in CG and CHG (10-14%) than CHH context (3- 5%) during embryonic development (embryonic cold or heat memory). Levels are lower in CG/CHH contexts after post-embryonic development (2-8%) but similar (1-5%) in CHH context (post-embryonic cold or heat memory). The number of unigenes containing at least 1 memorized temperature-DMC is higher (all contexts) for cold (4000-8000) than heat (2000- 5000) after both embryonic and post-embryonic development (Fig. 4C). In contrast, the percentage of development-DMCs during the embryonic phase at 23°C (E-SE vs. C-SE) that are memorized at the post-embryonic phase (C-SE vs. SAM, Fig. 5B, S3E) is much higher, reaching up to 45-50%, notably in CG/CHG contexts (post-embryonic developmental memory). Regarding promoters (Fig. 5B), no temperature memory could be detected while the post-embryonic developmental memory accounted for up to 90% of memorized DMCs (in CG). GO enrichment analysis for unigenes containing memorized temperature-DMCs (Fig. 6) revealed 6 major groups of GO terms found in 3 out of 4 situations, i.e., cold/heat embryonic (Fig. 6A) or post-embryonic memory (Fig. 6B): 4 groups related with i) heat (‘response to heat’, 2 times with the highest p-values), ii) UV (‘response to UV’, ‘pigment accumulation in response to UV light’), iii) the stress hormone jasmonic acid (‘response to jasmonic acid’, ‘jasmonic acid biosynthesis’), and iv) β-alanine (‘beta-alanine metabolism’) as a plant defense compound to withstand various stresses (Parthasarathy et al. 2019), and 2 metabolic groups related to v) ‘Glycolysis/Gluconeogenesis (once with the highest p-value), and vi) ‘starch and sucrose metabolism’.

**Figure 6:**
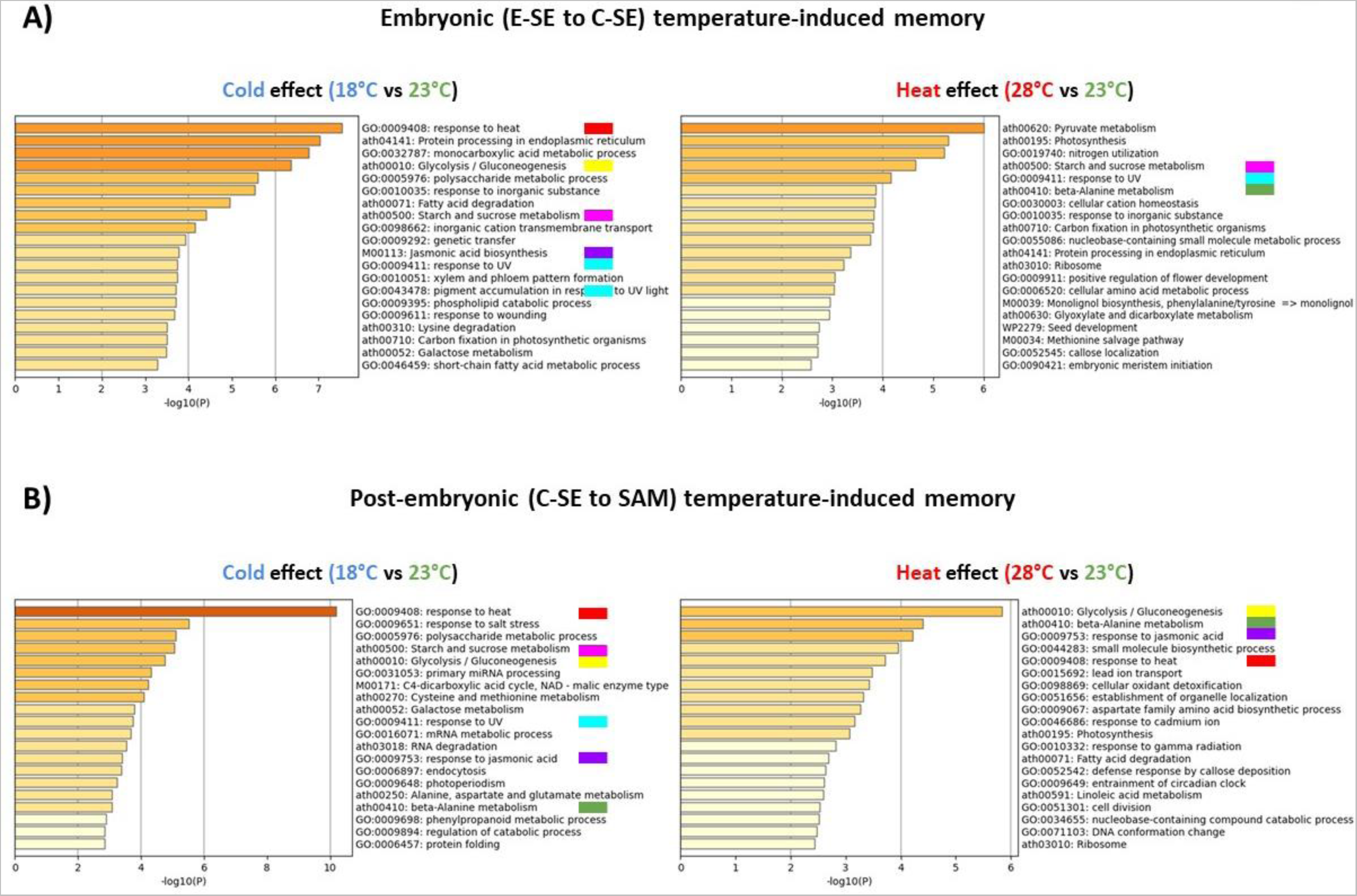
GO enrichment analysis of unigenes with memorized temperature-induced DMCs (memory) for all 3 contexts. **A.** Embryonic (E-SE to C-SE) development. **B.** Post-embryonic (C-SE to SAM) development. Most recurrent GO terms in at least 3 out of the 4 different cold (18/23°C) and heat (28/23°C) situations are highlighted by colored boxes: *Stress GO terms* related to response to heat (red), UV radiations (blue), jasmonic acid (purple), beta-alanine metabolism (green). *Metabolic GO terms* related to glycolysis /gluconeogenesis (yellow), starch and sucrose metabolism (pink).

### Identification of candidate genes under epigenetic control associated with temperature memory from embryonic to post-embryonic development

To identify the most significant candidate genes possibly involved in the establishment of a temperature memory, only those containing at least 5 memorized DMCs at the post-embryonic stage (C-SE to SAM) were selected. Excluding genes with no annotation, retrotransposons and rRNA genes, we identified a list of 10 candidate genes (Table 2). Post-embryonic cold memory genes correspond to a Subtilisin-like protein (cell division and embryo development), a Cystein- rich receptor-like protein (defense response), a Histone H3.1 protein (chromatin and regulation of cellular processes) and two proteins involved in the jasmonic acid (JA) pathway (DAD1-like lipase and Cytochrome P450). Post-embryonic heat memory genes match the same Cystein- rich receptor-like protein (defense response), but also a Heat shock protein (protection against heat stress), an Embryo defective MAPK kinase (embryo development, protein phosphorylation), an Arabinogalactan methylesterase (cell wall biogenesis), an ABC transporter family (developmental processes, adaptation), and an Insulinase family protein (embryo development).

**Table 2:**
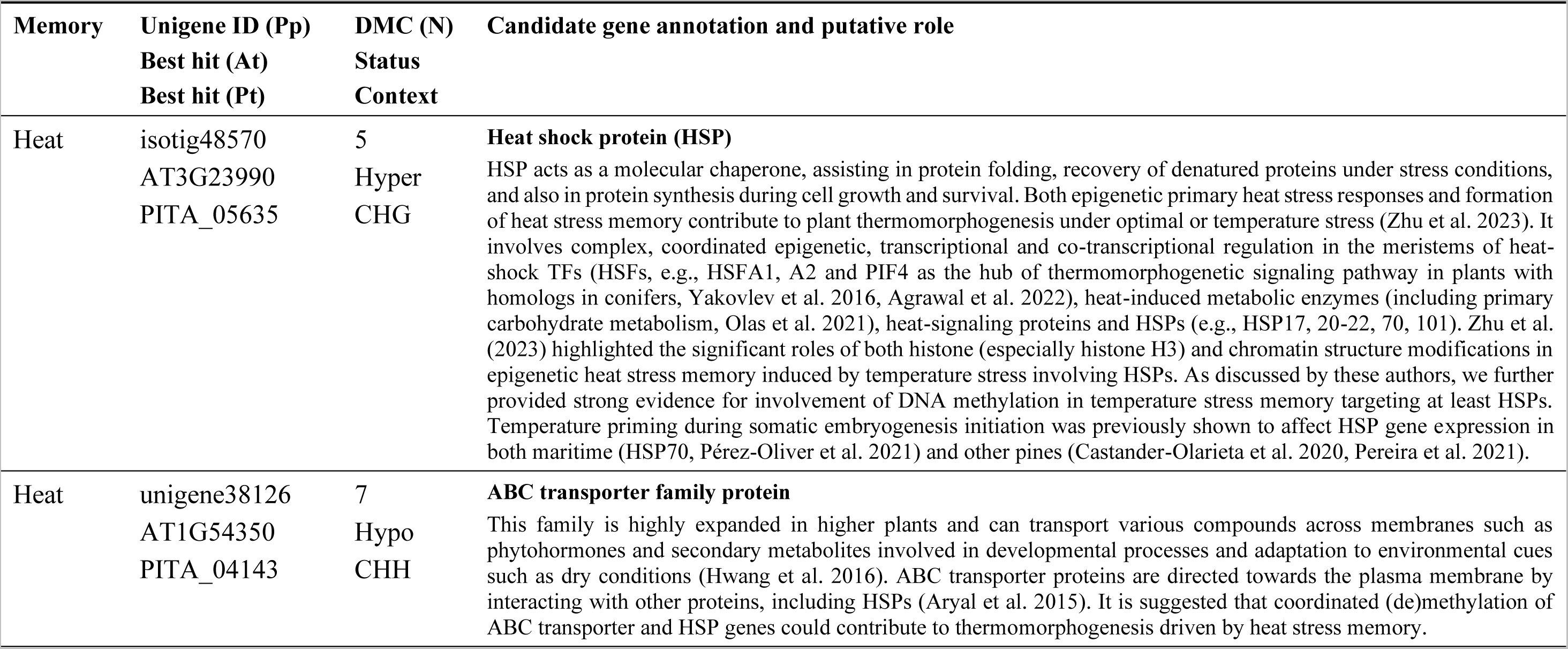
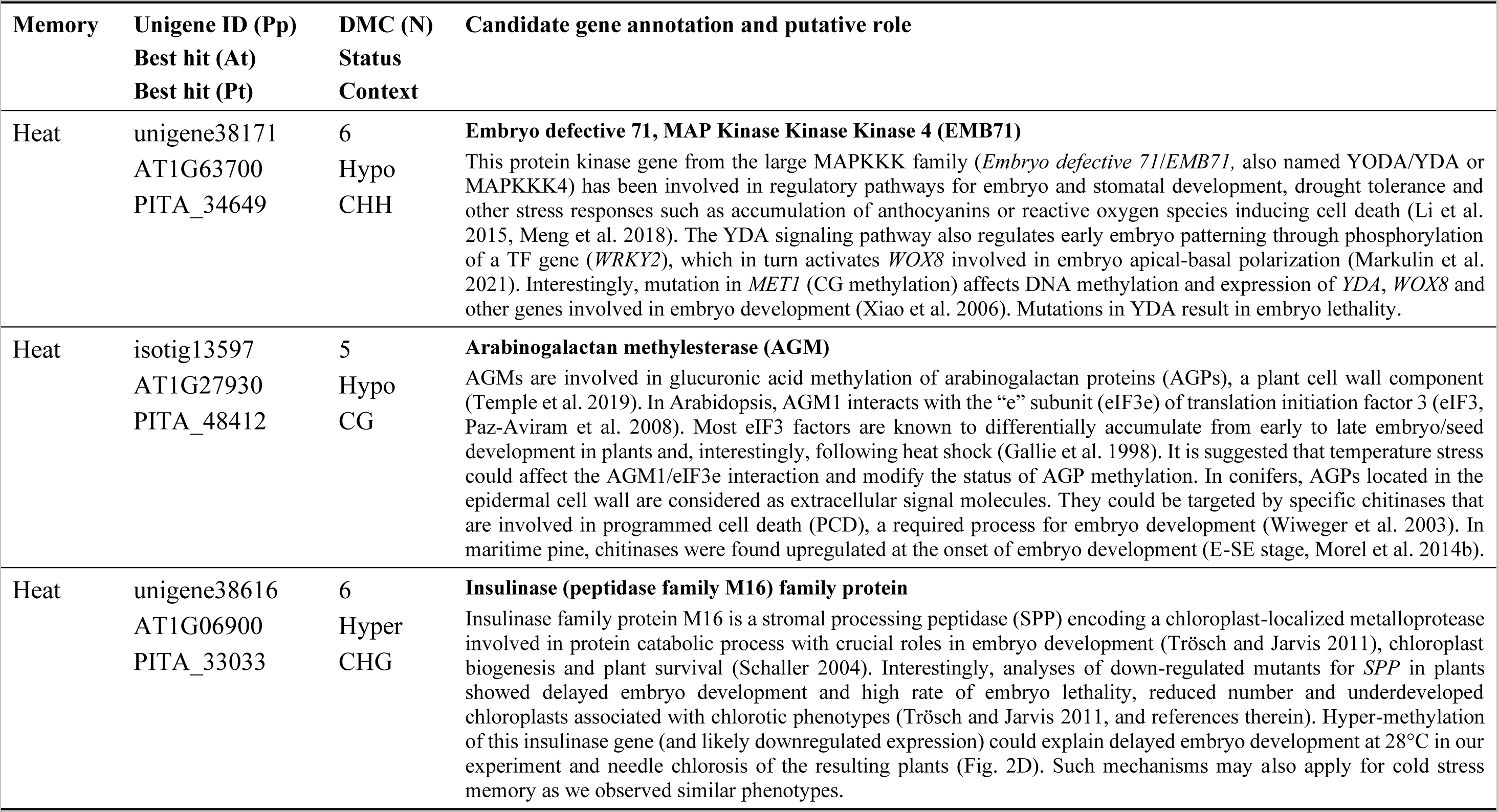
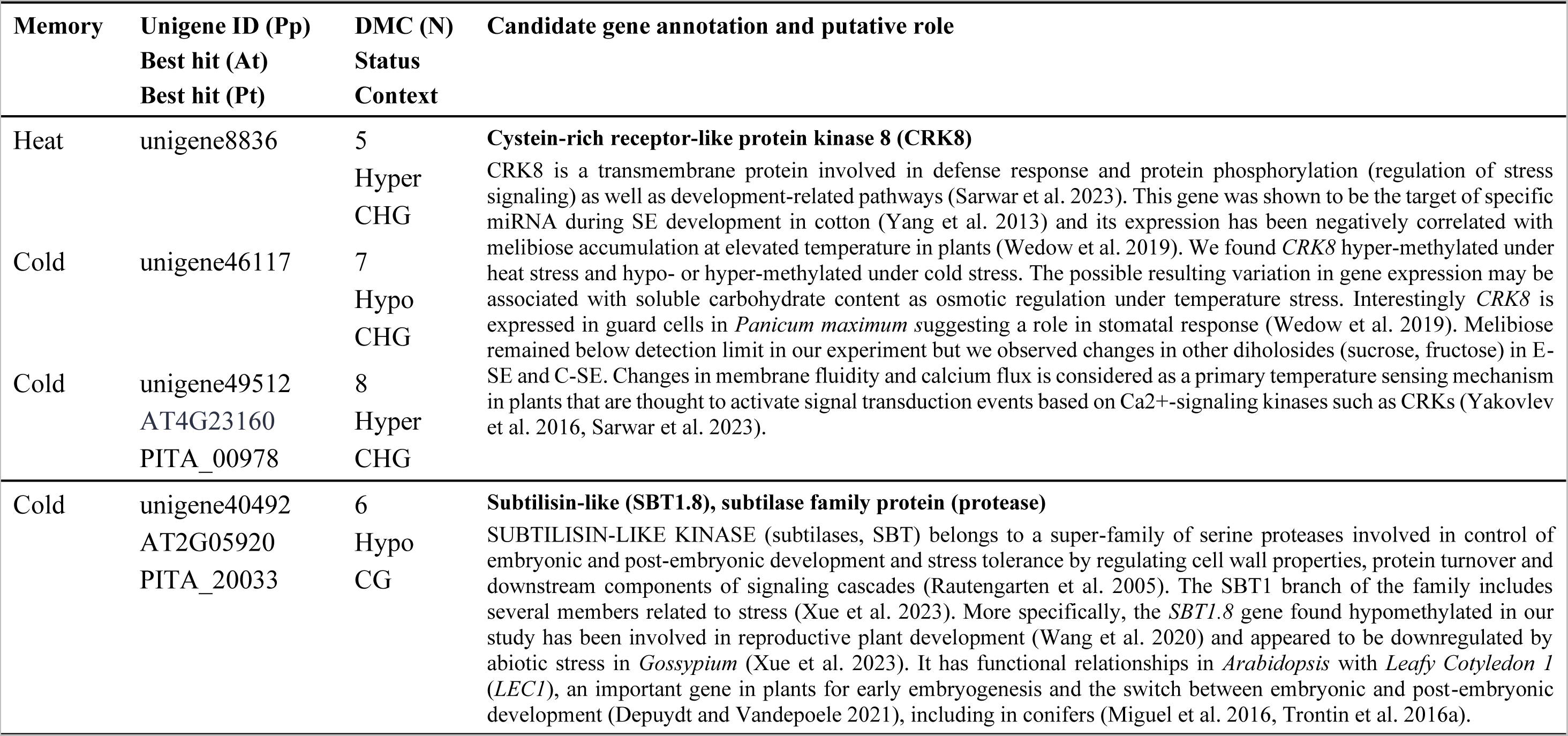
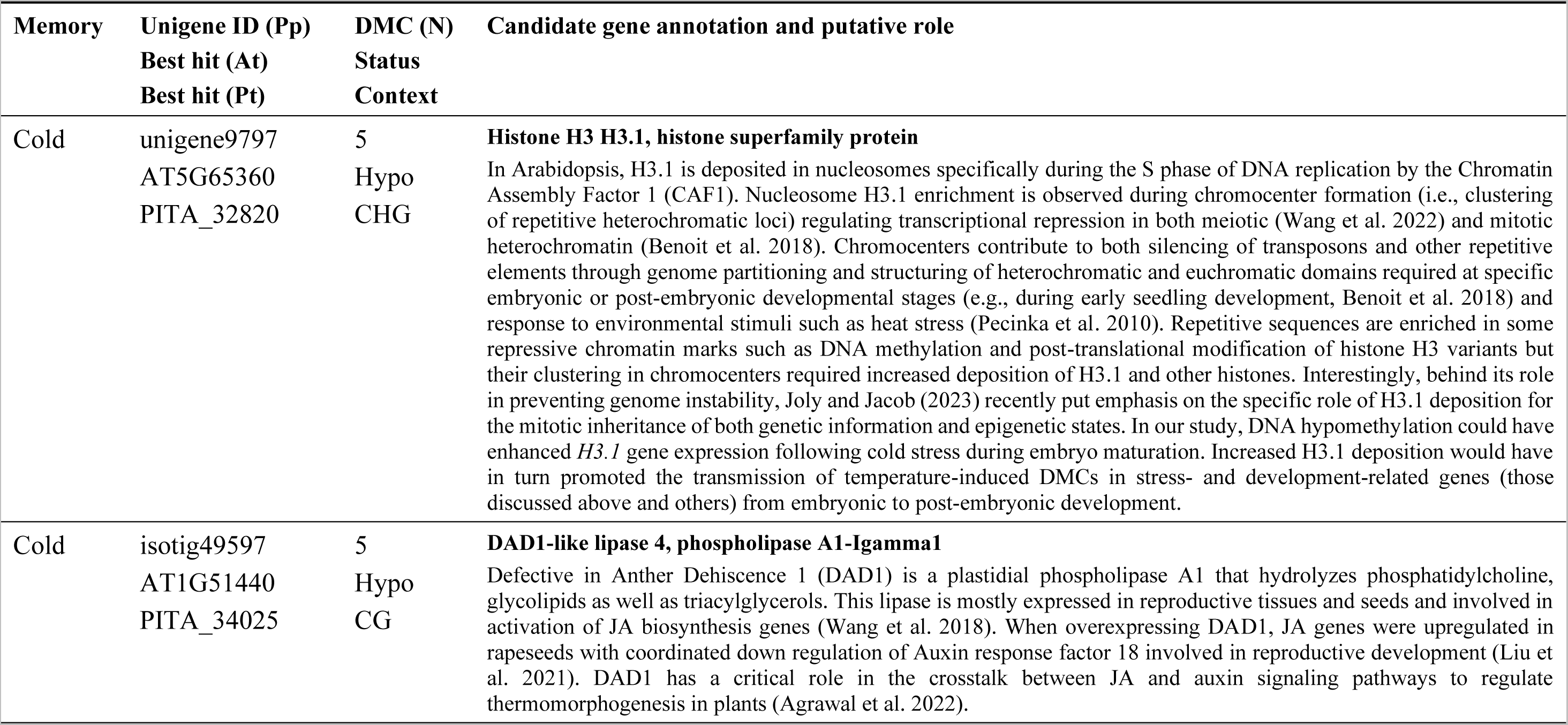
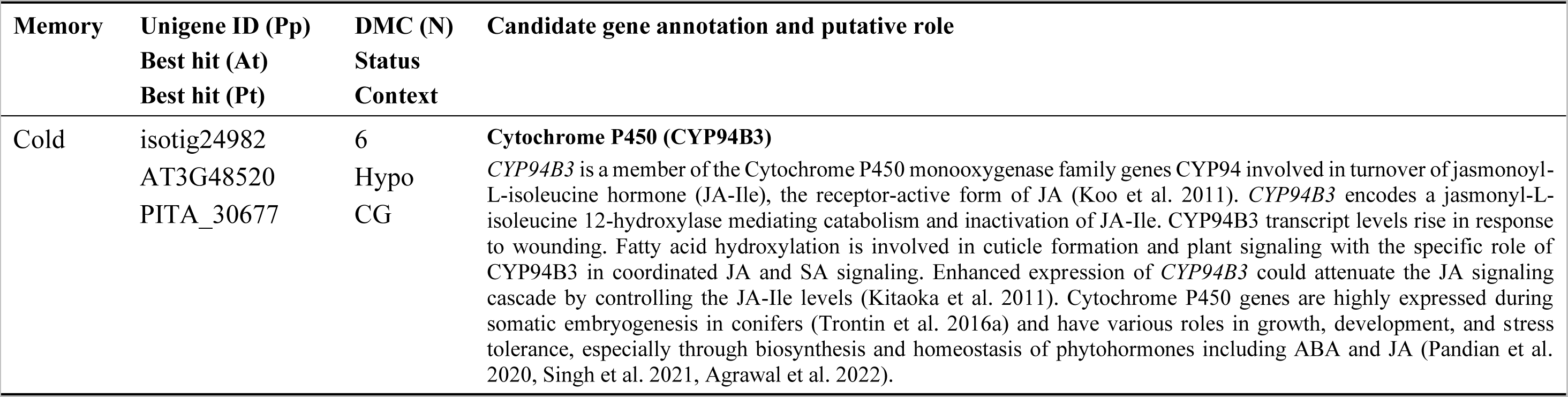
Best-annotated candidate genes for post-embryonic memory of temperature during embryo maturation in *Pinus pinaster*. Unigene ID in *Pinus pinaster* (Pp) and best corresponding hits in *Arabidopsis thaliana* (At) and *Pinus taeda* (Pt) are given for candidate genes containing at least 5 differentially-methylated cytosines (DMC) with consistent hyper- or hypo-methylation status in a given methylation context (CG, CHG, or CHH; H = A, C, or T). DMCs established in C-SE and memorized in the plant SAM in response to heat (28 vs. 23°C) or cold (18 vs. 23°C).

## DISCUSSION

### The methylome landscape of *Pinus pinaste*r somatic embryo and tree

Our SE material (Fig. S2B) showed high CG (65-70%) and CHG (55-60%) methylation levels compared to the CHH context (10-20%). This observation is consistent with gymnosperm data from needles (ca. 75/69/1-2%, respectively) and embryogenic tissue (ca. 65/60/3-4%, *Picea abies*, Ausin et al. 2016), shoots (ca. 88/82/2%, *Pinus tabuliformis*, Niu et al. 2022), meristems, and young leaves (ca. 78/76/36%, *Welwitschia mirabilis*, Wan et al. 2021). These values support specific roles of CHG/CHH methylation in conifers (Niederhuth et al. 2016, Ausin et al. 2016, Li et al. 2023).

We do not confirm that SE material has abnormal CG and CHG methylation patterns (Ausin et al. 2016), but rather specific ones associated with developmental stages (Fig. S4). Compared to other gymnosperm tissues, gene and promoter methylation appeared to be similarly high for C-SE in CG/CHG contexts (up to 70-75%, Fig. S4) but lower for E-SE and SAM. We confirm the high CHH methylation reported by Ausin et al. (2016), but it could be primarily related to development (Fig. S4) as CHH methylation was much higher in C-SE (up to 10-15% in genes, 20-35% in promoters) compared to E-SE and SAM (5-10% in genes, 10-20% in promoters).

We found higher CG (70 vs. 65%), similar CHG (ca. 55%) and lower CHH (10 vs. 20%) methylation in genes compared to promoters (Fig. S2B). In *P. abies*, Ausin et al. (2016) reported lower gene methylation in all 3 contexts compared to upstream regions. Our results are more aligned with data reported in *P. tabuliformis* (Li et al. 2023) when considering genes with transposable elements (TE, Niu et al. 2022). In this species, genes without TE insertion had much lower methylation levels than promoters in all 3 contexts. Interestingly, gene models had similar (CHH) or lower CG (60%) and CHG (45%) methylation than promoters in our study (Fig. S2B, S3). The regulatory role of gene and promoter methylation has been widely evidenced in angiosperms but was controversial in gymnosperms (Ausin et al. 2016). However, based on a high-quality genome assembly and annotation in *P. tabuliformis*, DNA methylation has been convincingly shown to be negatively correlated with gene expression (Li et al. 2023), especially when targeting exons and gene regions located downstream transcription end site (in all 3 contexts), but also promoters in a more limited way (only in CG context).

### The large developmental remodeling of the embryo methylome and epigenetic memory

Characteristic (epi)genomic changes in gene expression at key SE/ZE development stages have been reported in both angiosperms (Narsai et al. 2017, Ji et al. 2019, Chen et al. 2020) and conifers (Vestman et al. 2011, Trontin et al. 2016a). In maritime pine, de Vega-Bartol et al. (2013) reported large sets of differentially expressed genes between stages with evidence for transcriptional regulation by transcription factors (TFs) and epigenetic control, including DNA methylation, chromatin remodeling and non-coding sRNAs. Accordingly, DNA methylation (Fig. S2A) was high in C-SE (29%) compared to E-SE (15%) or plant SAM (25%) and hierarchical clustering (Fig. S2C) further pointed to a specific C-SE pattern, most notably in CHH/CHG contexts of promoters (Fig. S2C, S3B). Profound methylation rearrangements are suggested with an increase during the E-SE/C-SE transition followed by a decrease from embryonic to post-embryonic phase. Such changes have been reported in conifers with either similar (increased in C-SE from ca. 12 to 15-20%, Gao et al. 2022a) or opposite pattern (decreased in C-SE from ca. 61 to 53%, Teyssier et al. 2014).

Our methylome analysis revealed that genes exhibited more variation of DNA methylation than promoters in response to development. While similar in the CHH context (1-5%), SMP sites in development-induced DMC reached 6-11% (CG context) and 7-10% (CHG) in genes (Fig. 3C, S3D) as compared to less than 0.1-1.3% in promoters, respectively (Fig. 3D). This might be due to increased plasticity in gene bodies (where small changes have little effects) compared to promoters. Development-DMCs were found mostly hypo-methylated in genes and even more in promoters, notably for CHH and CHG contexts (Fig. S7, Table 1). Accordingly, we found about 25,000 unigenes (12%) containing embryonic and post-embryonic development-DMCs in all 3 contexts, and 4,000-7,000 promoters (8-14%) in the CHH context but less than 2,000 (4%) in the CHG or 200 (0.4%) in the CG context (Fig. 4B). The major finding that significant development-DMCs (Fig. S6C) are mainly hypo-methylated in all 3 contexts and rather prevalent in multiple genes is consistent with large developmental reprogramming during both the embryonic and post-embryonic phase towards active expression of large, selected gene pools (Li et al. 2023).

Development-induced DMCs during the embryonic phase can be transmitted to the post-embryonic phase (Fig. 5B, S3E) at a high rate in both genes (25-50%) and promoters (18-90%), especially in CG (50-90%) and CHG contexts (50-60%). Such a development-induced epigenetic memory was retained despite the likely massive epigenetic reprogramming during germination (Narsai et al. 2017, Tao et al. 2017, Markulin et al. 2021). This memory is likely to relate to genes whose level of expression established during embryogenesis is critical at least during the juvenile tree phase. We selected 2,500-8,500 genes and 50-1600 promoters containing at least 5 development-DMCs as a stringent criterion (Table 1, Fig. S8B). Main molecular functions of filtered DMCs for embryonic (Mat. S3) and post-embryonic effects (Fig. S9, Mat. S3) were mostly related to germination and early tree growth, as expected. Temperature and heat stress responses were also revealed in both genes and promoters, emphasizing that development is under strong temperature control for both acclimation and tolerance responses in plants (Zhu et al. 2023).

### Temperature-induced remodeling of the embryo methylome

Temperature is a critical factor affecting embryogenesis, plant growth, and development. We detected a temperature effect on SE hypocotyl size (Fig. S1A) that has been associated with thermomorphogenesis mechanisms involving major TFs (Perella et al. 2022, Zhu et al. 2023). Similar temperature-induced morphological changes were reported in pine (do Nascimento et al. 2020) and we further expanded the observed alterations to cotyledon number. We confirmed data in maritime and radiata pines (Sales et al. 2022, Moncaleán et al. 2018) showing faster growth (Fig. 2E) and/or differences in physiology (Fig. 2D) of somatic seedlings after SE maturation at 23°C compared to 18°C or 28°C. Temperature memory effects on early plant growth were also reported in *P. abies* (Kvaalen and Johnsen 2008) but, as in maritime pine (Fig. S1E), they did not appear to last for long (2 years) unlike phenological traits (at least 6 years, Skrøppa 2022). Both stability and instability of stress memory for different traits could be consistent with an epigenetic determinism.

We did not detect global changes in DNA methylation levels (Fig. S2A, 3C) in response to maturation temperature as reported in radiata pine (do Nascimento et al. 2022). In this species, significant variations were found in somatic seedlings by Castander-Olarieta et al. (2020), but following flash temperature treatments at the SE initiation step. Temperature effect could be more pronounced when applied at the early stages of somatic embryogenesis. However, no differences could be observed by Pereira et al. (2021) in Aleppo pine with a similar experimental design.

Despite a fairly stable global DNA methylation, we identified thousands of unigenes (ca. 25,000, 12%) in E-SE, C-SE, and SAM containing temperature-induced DMCs in all 3 methylation contexts (Fig. 4A). Both cold and heat stresses induced similar intensive responses in genes. In contrast, DMCs were found in less than 200 promoters (0.4%). In plants, gene methylation could be a key determinant of environmental stress response regardless of regulatory signals located in the promoter region such as TF binding sites (Aceituno et al. 2008). Regulation of gene expression could be restrained by DNA methylation, especially when occurring in gene exons as shown in pine (Li et al. 2023).

As for developmental effects, both genes and promoters containing temperature-induced DMCs showed an enrichment of molecular functions related to metabolisms and response to temperature stress (Fig. S9, Mat. S3). Similarly, we identified mainly in genes molecular functions associated with embryonic development and post-embryonic growth. Temperature sensing could involve a defense response through the jasmonic acid pathway as several related molecular functions have been identified in genes for both cold and heat stress.

### Epigenetic memory of temperature sensed during embryo maturation in 2-year-old regenerated somatic plants

Temperature-induced DMCs can be transmitted from embryonic to post-embryonic phases but, in contrast to development-induced DMCs, at a quite low rate (1-14% vs. 18-90%) and only in genes (Fig. 5B, S3E). We conclude that DNA methylation of the gene rather than the promoter may contribute to the establishment of an epigenetic memory of maturation temperature in maritime pine because it might be more permissive and less tightly controlled. Promoter methylation may be more related to a primary response to temperature involving transient TF binding rather than heat stress memory, which would require more persistent binding (Zhu et al. 2023). Our results are in line with transcriptional profiling of epigenetic regulators during SE development in Norway spruce (Yakovlev et al. 2016) showing that most genes involved in DNA (de)methylation are differentially expressed according to temperature and therefore predicted to be involved in the formation of an epigenetic memory. We significantly expanded upon this finding by demonstrating not only that methylation patterns are established in response to temperature during embryogenesis, but also that they are partially retained in the plant SAM. Temperature-DMCs are memorized at a lower frequency and preferentially in genes compared to development-DMCs, suggesting that DNA methylation contributes to temperature stress memory formation by targeting a different subset of genes and/or genes that are not essential for proper embryo and seedling development. The later hypothesis is supported by our data showing no temperature effect on SE viability and germination rates (Fig. S1C). Yakovlev et al. (2016) discussed that genes involved in embryo development could be weakly responsive to temperature. However, we found good support for molecular functions related to thermosensing of both genes and promoters containing development-DMCs (Fig. S9, Mat. S3). Differential responses to temperature of genes involved in embryogenesis and plant development could partly explain why more memorized temperature-DMCs were observed at the embryonic phase (3-14%) compared to the post-embryonic phase (1-8%) in some contexts (CG, CHG).

We found more genes with memorized temperature-DMCs for cold (4000-8000) than for heat (2000-5000) (Fig. 4C). Compared to the reference temperature, cold therefore was a stronger stimulus than heat, emphasizing also that the response could be non-linear. Significant phenotypic variations were indeed observed after maturation at 18°C (Fig. 2, S1) in E-SE (reduced proliferation and glucose content, increased starch content), C-SE (delayed development, reduced yield, fresh mass, hypocotyl size, cotyledons number, and starch content), and plant (chlorosis, reduced growth). Maturation at 28°C had comparatively lower effects.

Molecular functions associated with genes containing memorized temperature-induced DMCs (Fig. 6) include defense mechanisms (response to heat, UV, or JA, β-alanine metabolism) and carbon metabolisms (glycolysis, gluconeogenesis, starch and sucrose). In accordance with phenotypic data (Fig. 2, Fig. S1), they were more consistently observed for cold stress. This result emphasized DNA (de)methylation as a regulator of the expression of genes involved in the JA pathway for sensing temperature, not only as a primary perception, but also for stress memory establishment. JA signaling has important roles in plant thermomorphogenesis (Clarke et al. 2009, de Ollas et al. 2015, Khan et al. 2022, Agrawal et al. 2022) as well as in embryo and plant development (Elhiti et al. 2013, Wang et al. 2018). It operates through a complex crosstalk with other phytohormone signaling pathways (Maury et al. 2019), especially salicylic acid, ABA, ethylene (stress) and auxin (development). Activation of the JA pathway could be coordinated with regulatory networks such as epigenetic-induced response to heat (Zhu et al. 2023), production of β-alanine as a generic stress response (Parthasarathy et al. 2019), and modulation of primary carbon and carbohydrate metabolisms as key regulators of embryo development (Trontin et al. 2016a) contributing to heat-stress memory (Olas et al. 2021). Accordingly, we found temperature-related changes in glucose, sucrose and starch in both E-SE and C-SE (Fig. S1B).

Ten robust candidate genes for epigenetic temperature sensing and memory formation in maritime pine were identified (Table 2). In good concordance with the GO enrichment analyses (Fig. 6, Fig. S9, Mat. S3), they mostly encode proteins involved in defense responses and adaptation and/or embryo development, and also chromatin regulation (see Table 2 for putative role of each gene). Hyper- or hypo-methylation of *Cystein-rich receptor-like protein kinase* gene (*CRK8*) could support both cold and heat memory. Hyper-methylation of *Heat shock protein* gene (*HSP*) and hypo-methylation of *ABC transporter family* gene were related with heat stress memory. The JA pathway role in cold stress memory is emphasized by hypo- methylation of *DAD1-like lipase 4* and *Cytochrome P450* (*CYP94B3*). Regarding development-related genes, cold stress memory is supported by hypo-methylation of one *Subtilisin-like kinase* gene (*SBT1.8*). Heat stress memory involves hypo-methylation of *Embryo defective 71 kinase* (*EMB71*) and *Arabinogalactan methylesterase* genes, and hyper-methylation of *Insulinase family protein M16* gene. Interestingly, cold stress memory was also supported by hypo-methylation of the histone *H3.1* gene involved in the regulation of cellular processes through chromatin rearrangement.

### Epigenetic control of temperature memory in trees: prospects

DNA methylation (5mC) is currently the epigenetic modification, which can most easily be monitored at both gene and genome levels during memory formation. In the context of climate change, the possibility for early temperature priming during embryogenesis to stimulate plant phenotypic plasticity (e.g., increased tolerance to drought and heat stress) through epigenetic stress memory has great potential in plants, especially in trees with long life cycle and delayed reproductive maturity (Liu et al. 2021, 2022, Sterck et al. 2022, Zhu et al. 2023). If transgenerational stress memory currently appears to be out of reach for trees in an acceptable time frame, intergenerational stress memory can be considered for (epi)genetic breeding applications, including in conifers (Liu and He 2020). Somatic embryogenesis is primarily a promising way to multiply zygotic embryos in a context of seed shortage. Our results and others in conifers (Kvaalen and Johnsen 2008, Moncaleán et al. 2018, Arrillaga et al. 2019, do Nascimento et al. 2020, Sales et al. 2022) showed that temperature variation during early embryogenesis can affect both SE yield and quality and is therefore a way of refining the process towards production of vigorous seedlings of forestry standard, a current limitation of somatic embryogenesis in maritime pine and other conifers (Trontin et al. 2016b, Lelu-Walter et al. 2016). DNA methylation marks associated with optimal SE development as a function of environmental conditions from initiation to maturation (temperature and other abiotic factors: water availability, hormonal balance, nutrition, etc.) could provide new tools for easy monitoring of embryogenic cultures and embryo quality. As a key clonal process to access various technologies in conifers (e.g., cryopreservation, genomic selection, genetic engineering, Klimaszewska et al. 2016), somatic embryogenesis further provides a convenient tool for early priming with abiotic stress in controlled conditions. Our data suggest that temperature-induced memory can be used to modulate early growth of somatic seedlings. Even if temporary, this effect can have important applications in forestry because initial seedling vigor is recognized as a crucial factor for competing with weeds during the first growing season in the field, especially in maritime pine (Trontin et al. 2016b). Interestingly, temperature memory of somatic seedlings can apparently be used to modulate various traits of high interest for plantation forestry in conifers such as plant developmental phenology, tolerance to heat and drought stress (Kvaalen and Johnsen 2008, Castander-Olarieta et al. 2020, Pérez-Oliver et al. 2021, 2023, do Nascimento et al. 2022, Sales et al. 2022). Here again, DNA methylation marks could be useful (in conjunction with genetic markers) for managing the production of well-adapted (epi)genetic resources.

## Supporting information

Supplementary Material 1 - Biological and biochemical data

Supplementary Material 2 - Designed probes for sequence capture bisulfite (SeqCapBis design, Roche)

Supplementary Material 3 - Gene Ontology (GO) enrichment analysis (Metascape)

## Acknowledgments

M.D.S. received PhD grants from the French Ministry of Higher Education and Research (MESR). This work was supported by the Centre-Val de Loire Region (France) through the INTEMPERIES project (grant n°2014 00094511 to INRAE, Coord. M.A.L.W.). Scientific and technical support was provided by the epigenomic environmental platform (P2E) of IHPE (http://ihpe.univ-perp.fr/plateforme-epigenetique/, Perpignan, France) and by the shared FCBA-INRAE XYLOBIOTECH platform (grant n°ANR-10- EQPX-16, Cestas, Orléans, France), a service from the IN-SYLVA France Research Infrastructure (https://in-sylva-france.hub.inrae.fr/services/in-lab). This work was also supported by a grant overseen by the French National Research Agency (ANR) as part of the “Investissements d’Avenir” program (ANR-11-LABX-0002-01, Lab of Excellence ARBRE).

## Author contributions

S.M. coordinated the research. The plant experimental design was established by J.F.T, M.A.L.W. and S.M. Somatic embryo culture and measurements were performed by I.R. and J.F.T. Somatic plant growth and phenotypic assessments were performed by F.C. and J.F.T. Biochemical analyses were performed by N.B., C.L.M., and C.T. Statistics were performed by J.F.T. (biological data), C.T. (biochemical data), and S.M. (DNA methylation data). DNA extractions and capture were done by A.D. and M.D.S. under ROCHE supervision. HPLC analysis was performed by A.D., M.D.S., and S.M. Sequencing was performed by J.T. and C.D. Design was prepared by A.G. and S.M. Methylome data analysis was done by S.M. and C.C. Gene Ontology analysis was performed by I.M. and S.M. The draft manuscript was conceived and written by J.F.T, C.M., and S.M., and further revised by M.A.L.W. and C.M. All authors approved the final version of the manuscript.

## Data availability

The data that support the findings of this study are openly available. The Sequence Capture Bisulfite data have been deposited in NCBI at https://www.ncbi.nlm.nih.gov/bioproject/PRJNA874210, SRA: SRR21374316, SRR21374315, SRR21374314, SRR21374313, SRR21374312, SRR21374311, SRR21374310, SRR21374309, SRR21374317. Phenotypic data can be found in Mat. S1. Information about the sequence capture design (Roche) and all data from the Metascape analysis are shown in Mat. S2 and S3, respectively, and are available on *data.gouv.fr*, the open platform for French public data (doi: 10.57745/PNGW7G).

