## Supplementary Tables and Figures for "Epigenetic memory of temperature sensed during somatic embryo maturation in 2-year-old maritime pine trees"

### Short title: Epigenetic memory in pine trees

J.-F. Trontin[1§$], M.D. Sow[2£], A. Delaunay[2], I. Modesto[3], C. Teyssier[4], I. Reymond[1], F. Canlet[5], N. Boizot[4], C. Le Metté[4], A. Gibert[2], C. Chaparro[6], C. Daviaud[7], J. Tost[7], C. Miguel[3], M.-A. Lelu-Walter[4] and S. Maury[2$]

[1]BioForBois, FCBA, Pôle Industrie Bois & Construction, Cestas, 33610, France.
[2]P2e, Université d'Orléans, INRAE, EA 1207 USC 1328, 45067 Orléans, France.
[3]Biosystems and Integrative Sciences Institute, Faculdade de Ciências, Universidade de Lisboa, 1749-016 Lisboa, Portugal.
[4]BioForA, INRAE, ONF, UMR 0588, 45075 Orléans, France.
[5]Sylviculture Avancée, FCBA, Pôle Ressources Forestières des Territoires, Cestas, 33610, France.
[6]IHPE, Université de Perpignan, UMR 5244, 66100, Perpignan, France.
[7]Laboratory for Epigenetics and Environment, Centre National de Recherche en Génomique Humaine, CEA - Institut de Biologie François Jacob, Université Paris Saclay, 91000 Evry, France.

[§]Present address: BEF, INRAE, UR 1138, 54280 Champenoux, France.
[£]Present address: GDEC, INRAE, UMR 1095, 63000 Clermont-Ferrand, France.

**Key message:** Developmental and temperature-induced changes in the methylome of maritime pine somatic embryos can be stably transmitted from the embryonic to the post-embryonic phase.

**Keywords:** *Pinus pinaster*, somatic embryogenesis, memory, epigenetics, development, heat/cold stress, DNA methylation, methylome, sequence capture bisulfite.

### Supporting Information

#### Supplementary Tables

**Table S1:** Yield in PN519 cotyledonary somatic embryos (no. embryos/g f.m.) harvested after 12-16 weeks maturation according to the temperature and initial density of the embryogenic cell inoculum.

| Temperature (°C) | Cell density (mg / filter) | Maturation time (weeks) | | |
|---|---|---|---|---|
| | | **12** | **14** | **16** |
| **18** | 100 | 5 | 16 | 28 |
| | 50 | 4 | 4 | 36 |
| **23** | 100 | 46 | 80 | 132 |
| | 50 | 76 | 148 | 236 |
| **28** | 100 | 14 | 34 | 36 |
| | 50 | 36 | 64 | 98 |

Note: To estimate whether temperature affects the phenology of embryo development, we assessed the evolution of C-SE yield after 12-, 14-, and 16-weeks maturation at two cell plating densities (50 and 100 mg/filter). C-SE yield was higher at reduced cell density compared to our standard, especially at 23 and 28°C. Considering as a reference C-SE yields obtained after maturation for 12 weeks at 23°C (46-76 SE/g), both lower (18°C) and higher temperature (28°C) significantly delayed embryo development. It takes 15-16 weeks at 28°C to obtain similar yields to the control (36-98 SE/g). At 18°C, even after 16 weeks, the yield remains lower (28-36 SE/g), and the necessary maturation time of C-SE was estimated in subsequent experiments to be 17-18 weeks.

**Table S2:** Plant survival (%) under greenhouse conditions recorded 5, 8 and 15 months after germination of PN519 cotyledonary somatic embryos matured at 18, 23 or 28°C.

| **Plant age (months)** | **Temperature (°C)** | | |
|---|---|---|---|
| | **18** | **23** | **28** |
| **5** | 86.5 | 87.5 | 84.4 |
| **8** | 85.4 | 83.3 | 82.3 |
| **15** | 81.2 | 72.9 | 80.2 |

**Table S3:** Mapping efficiency of Sequence Capture Bisulfite using BSMAP on the three genomic designs: (1) 866 gene models (Seoane-Zonjic et al. 2016), (2) 206,574 unigenes (Cañas et al. 2017), and (3) 51,749 promoters (Zimin et al. 2017). For each biological sample (E-SE, C-SE, SAM at 18, 23 or 28°C) and design, the coverage is defined as the average of times the genomic design is covered by uniquely mapped paired reads.

| Samples | Genomic design | Uniquely mapped paired reads | Coverage (x) |
|---|---|---|---|
| E-SE, 18°C | Gene models | 1,973,206 | 310 |
| | Unigenes | 22,365,351 | 60 |
| | Promoters | 8,034,927 | 19 |
| E-SE, 23°C | Gene models | 2,081,112 | 327 |
| | Unigenes | 26,243,503 | 71 |
| | Promoters | 10,015,375 | 23 |
| E-SE, 28°C | Gene models | 1,189,466 | 187 |
| | Unigenes | 18,247,471 | 49 |
| | Promoters | 8,420,159 | 20 |
| C-SE, 18°C | Gene models | 1,295,595 | 204 |
| | Unigenes | 19,112,420 | 52 |
| | Promoters | 7,713,466 | 18 |
| C-SE, 23°C | Gene models | 1,331,013 | 209 |
| | Unigenes | 19,110,567 | 52 |
| | Promoters | 8,271,150 | 19 |
| C-SE, 28°C | Gene models | 427,161 | 67 |
| | Unigenes | 12,531,317 | 34 |
| | Promoters | 6,244,470 | 15 |
| SAM, 18°C | Gene models | 1,800,410 | 283 |
| | Unigenes | 25,934,571 | 70 |
| | Promoters | 9,848,747 | 23 |
| SAM, 23°C | Gene models | 1,577,441 | 248 |
| | Unigenes | 21,395,009 | 58 |
| | Promoters | 7,897,401 | 18 |
| SAM, 28°C | Gene models | 1,778,631 | 279 |
| | Unigenes | 26,706,657 | 72 |
| | Promoters | 10,773,607 | 25 |

*Note*: E-SE: Early Somatic Embryo; C-SE: Cotyledonary Somatic Embryo; SAM: plant Shoot Apical Meristem.

### Supplementary Figures

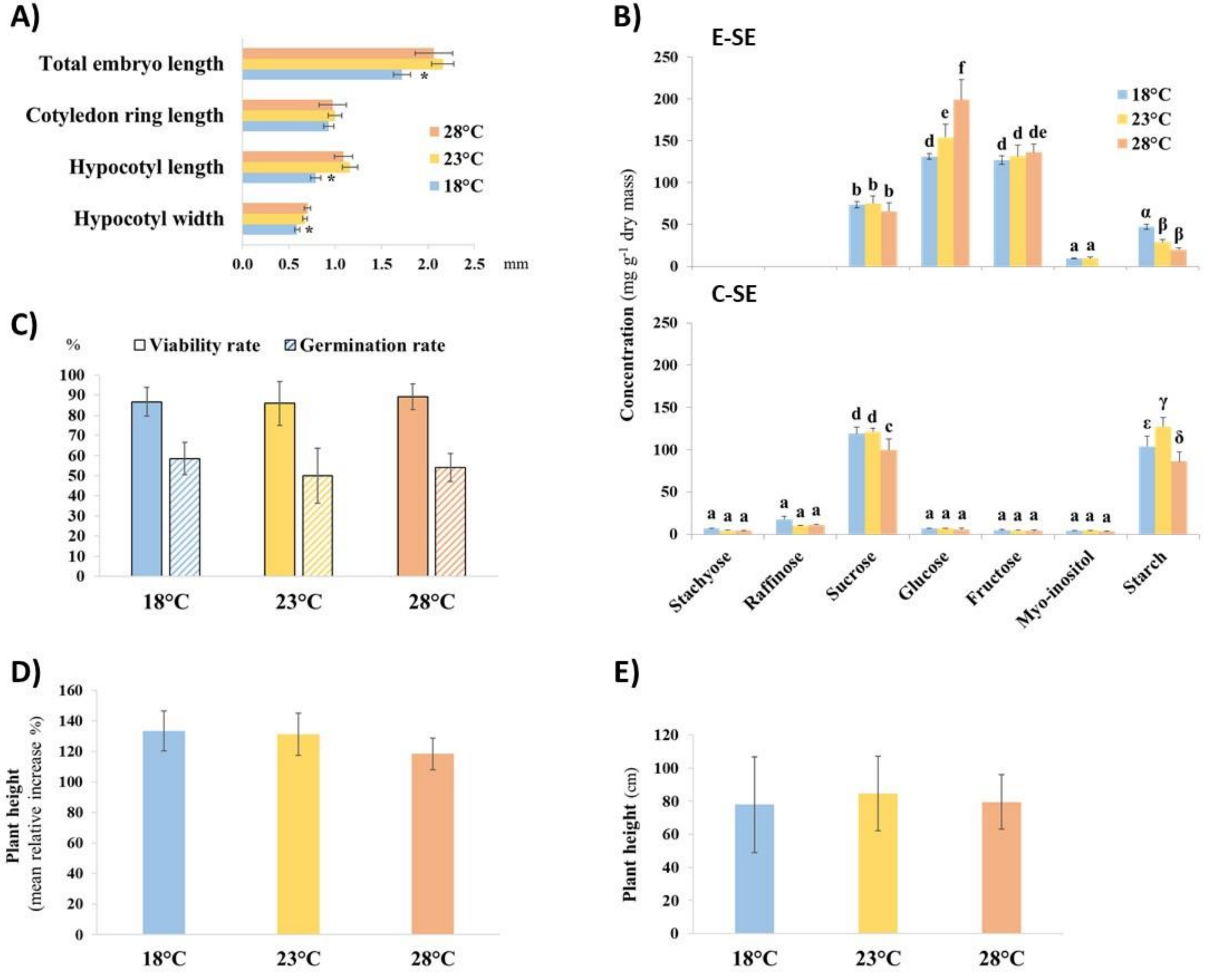


**Figure S1:** Effect of temperature during maturation (18, 23, 28°C) on embryonic and post-embryonic development of PN519 somatic embryos.

**A.** Mean size of cotyledonary somatic embryos (C-SE). Bars: 95% confidence limits. For each quantitative variable (total length, cotyledon ring length, hypocotyl length and width), significant differences are indicated by a star ($p < 0.05$).

**B.** Concentration in starch and detected soluble sugars in early (E-SE) and cotyledonary somatic embryos (C-SE). Bars: standard deviation. As the two types of data cannot be compared (because obtained by different assay methods) significant differences among temperature conditions and developmental stages are indicated by different latin (soluble sugars) or greek (starch) letters ($p < 0.05$).

**C.** Viability and germination rates of cotyledonary SE (C-SE). Bars: 95% confidence limits. For each variable, observed differences among temperature treatments are not significant.

**D.** Mean relative increase in plant height between 8 and 15 months. This increase essentially represents the annual spring flush. Bars: 95% confidence limits. Observed differences among temperature treatments are not significant.

**E.** Mean plant height at age 65 months (H65). Field growth. Bars: 95% confidence limits. No significant differences could be detected due to higher variability resulting from game damage.

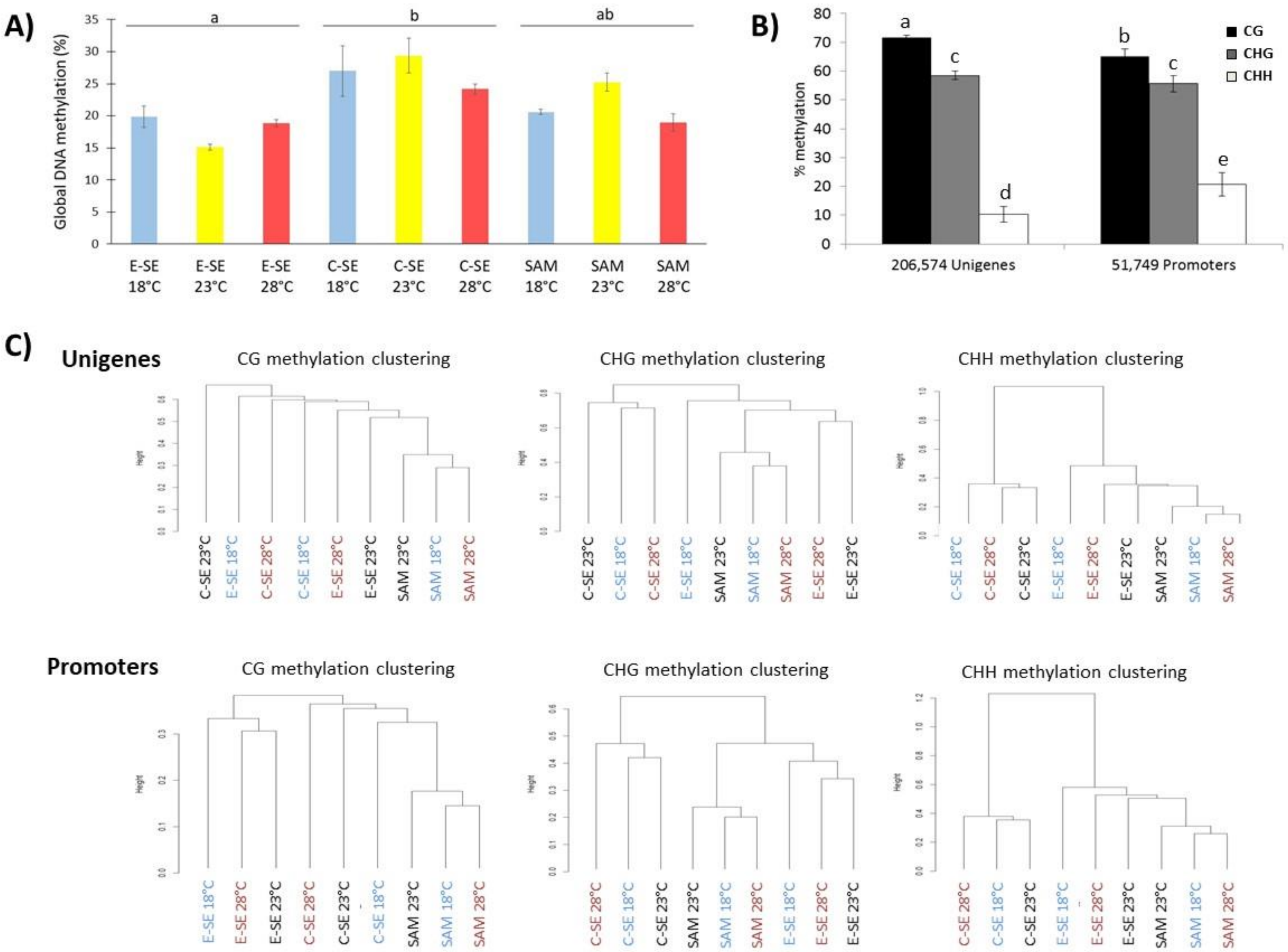


Figure S2: Global methylome analysis.

**A.** Global DNA methylation percentage (using HPLC) for each developmental stage (E-SE, C-SE, SAM) and temperature (18, 23, 28°C). Mean values are indicated with their standard errors (n = 3). Significant differences for means with ANOVA test are indicated by different letters.

**B.** Cytosine methylation percentage by context (CG, CHG, CHH) (using SeqCapBis) for unigenes and promoters. Mean values calculated from biological samples are indicated with their standard errors (n = 9) for both feature and cytosine context. Significant differences for means are indicated by distinct letters.

**C.** Hierarchical clustering of biological samples based on SMPs in the 3 cytosine contexts (CG, CHG, CHH) for unigenes (up) and promoters (down)

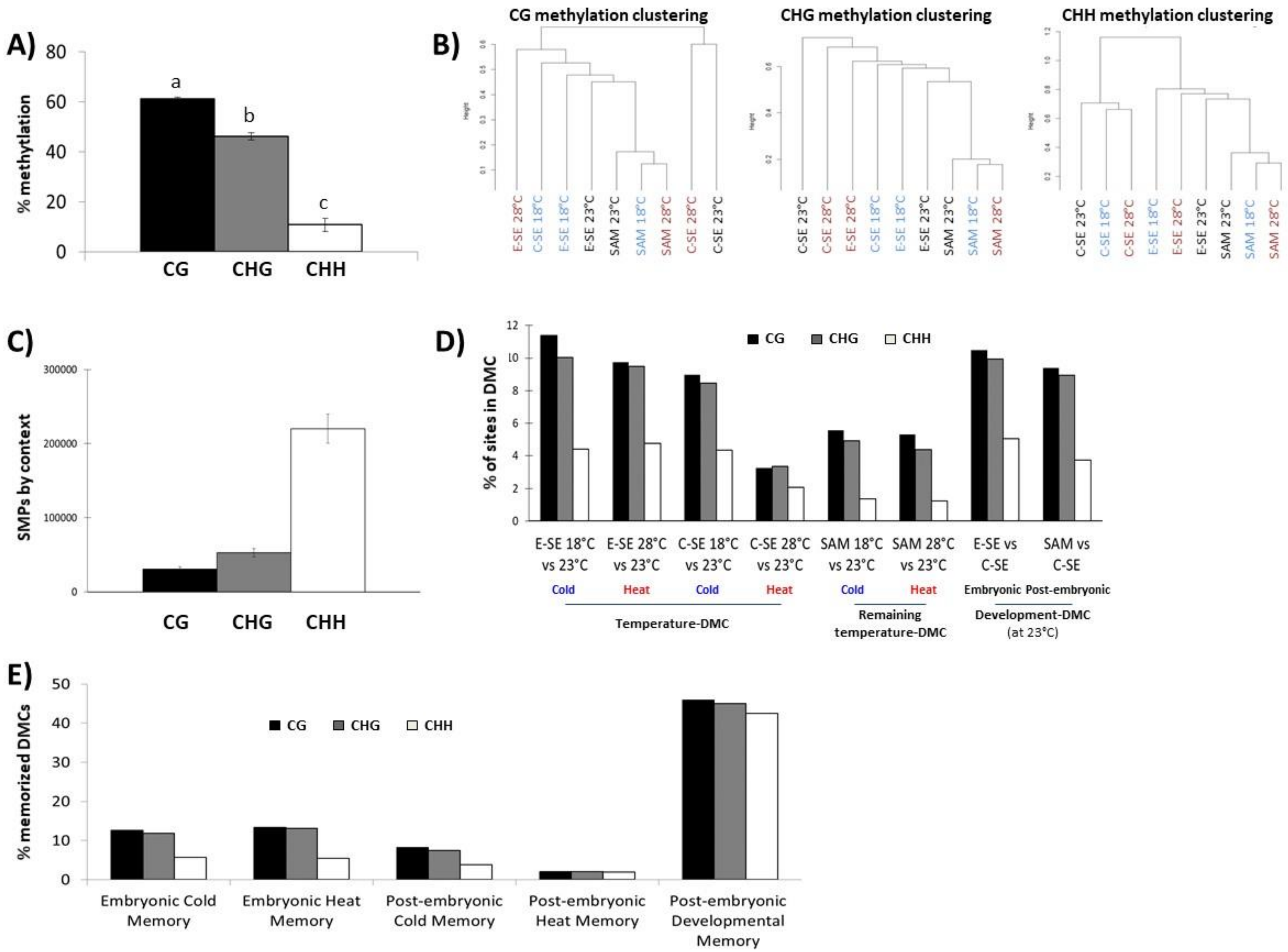


**Figure S3:** Methylome analysis of the 866 gene models in the 3 cytosine contexts (CG, CHG, CHH).

**A.** Cytosine methylation percentage (using SeqCapBis). Mean values for the biological samples with their standard error (n = 9). Significant differences for means are indicated by different letters.

**B.** Hierarchical clustering of biological samples based on SMPs.

**C.** Number of SMPs.

**D.** Percentages of SMP sites in significant DMC, i.e., with q-value < 0.01 and at least 25% methylation difference, compared to the total number of SMP sites analyzed for all *P. pinaster* samples. The different types of DMC (cold/heat temperature-, remaining temperature- and embryonic/post-embryonic development-DMC) are indicated (see Fig. S6).

**E.** Percentage (%) of memorized DMCs induced by temperature (cold: 18 vs. 23°C; heat: 28 vs. 23°C) or embryo development (C-SE vs. E-SE at 23°C). Post-embryonic developmental memory and both embryonic and post-embryonic temperature-induced (cold or heat) memories are shown.

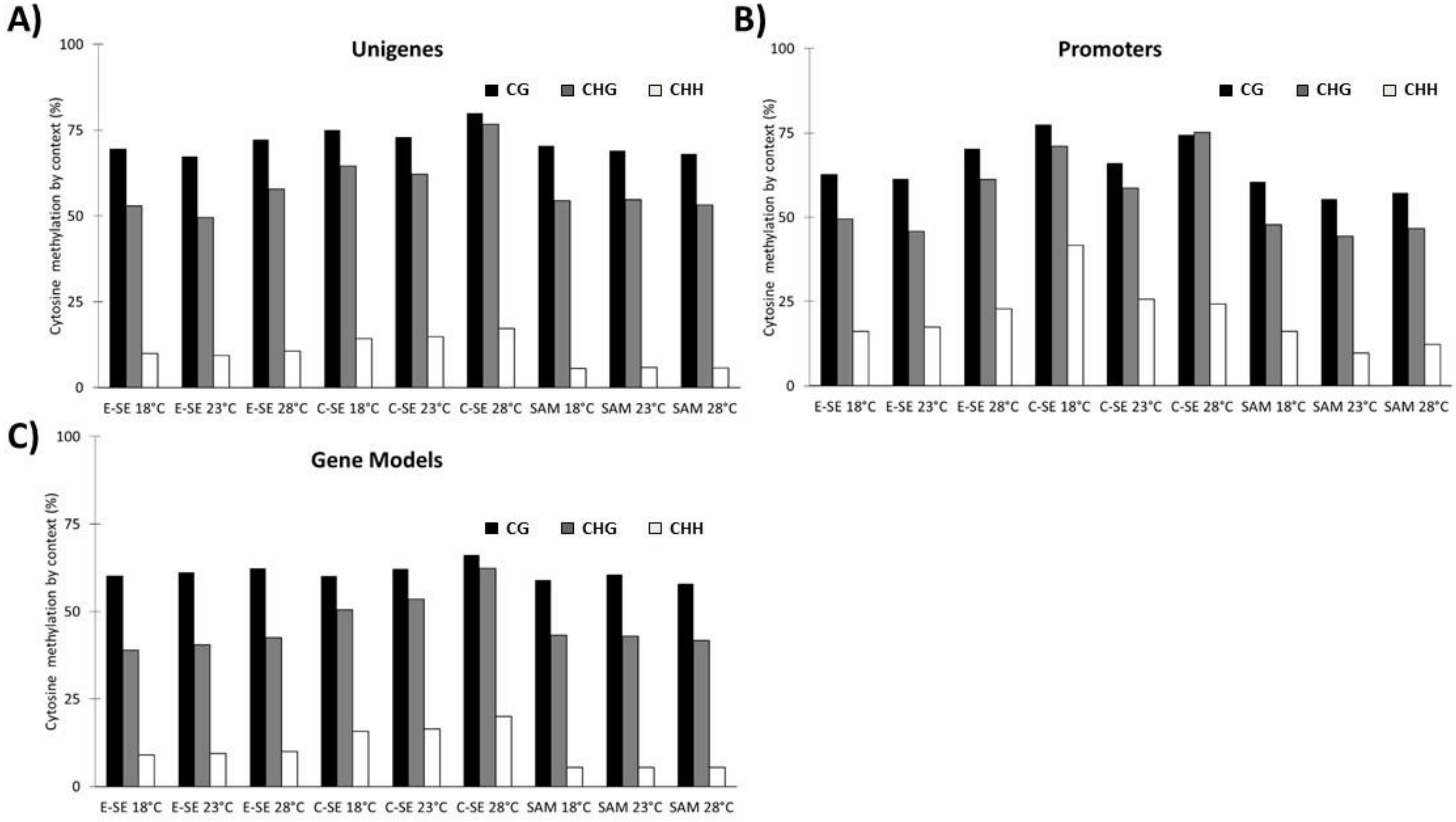


**Figure S4:** Cytosine methylation percentages by context (CG, CHG, CHH) (using SeqCapBis) in the 9 biological samples (E-SE, C-SE and SAM primed during maturation at 18, 23 or 28°C).

**A.** Unigenes; **B.** Promoters; **C.** Gene models**.**

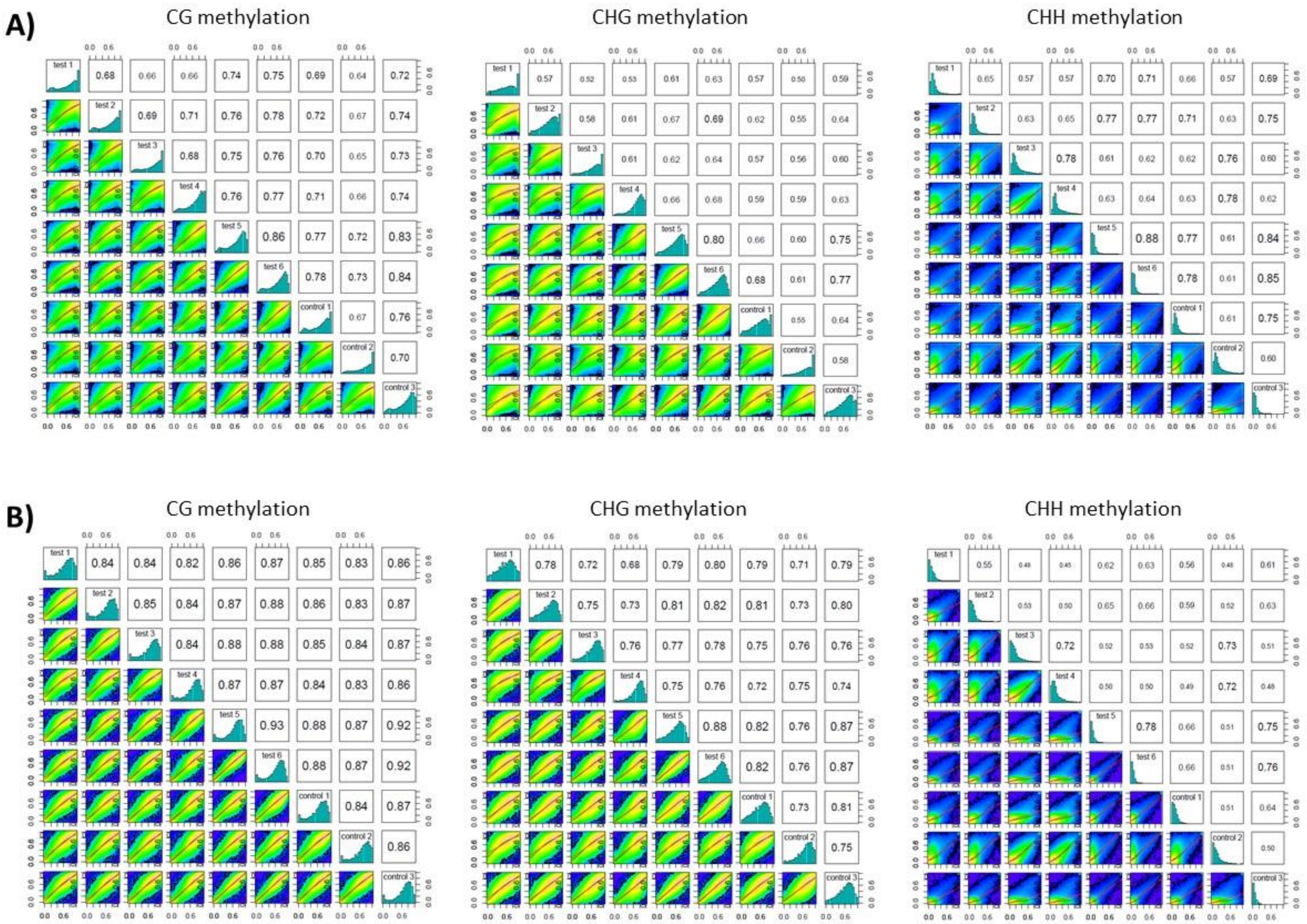


**Figure S5**: Correlations based on SMP values between the 9 *P. pinaster* samples considering the unigenes (**A**) or promoters (**B**) design by cytosine context (CG, CHG, CHH).

Scatter plots of correlation values for each pair of biological samples (E-SE, C-SE and SAM primed during maturation at 18, 23 or 28°C). Numbers on the upper right corner denote pairwise Pearson's correlation scores. SMP values histograms for each sample are on the diagonal. Data on the lower left corner are correlation curves among samples (the blue color indicates lower correlation than green and red). Test 1: E-SE 18°C, test 2: E-SE 28°C, test 3: C-SE 18°C, test 4: C-SE 28°C, test 5: SAM 18°C, test 6: SAM 28°C. Controls 1, 2 and 3 correspond to the reference temperature (23°C) for E-SE, C-SE and SAM, respectively.

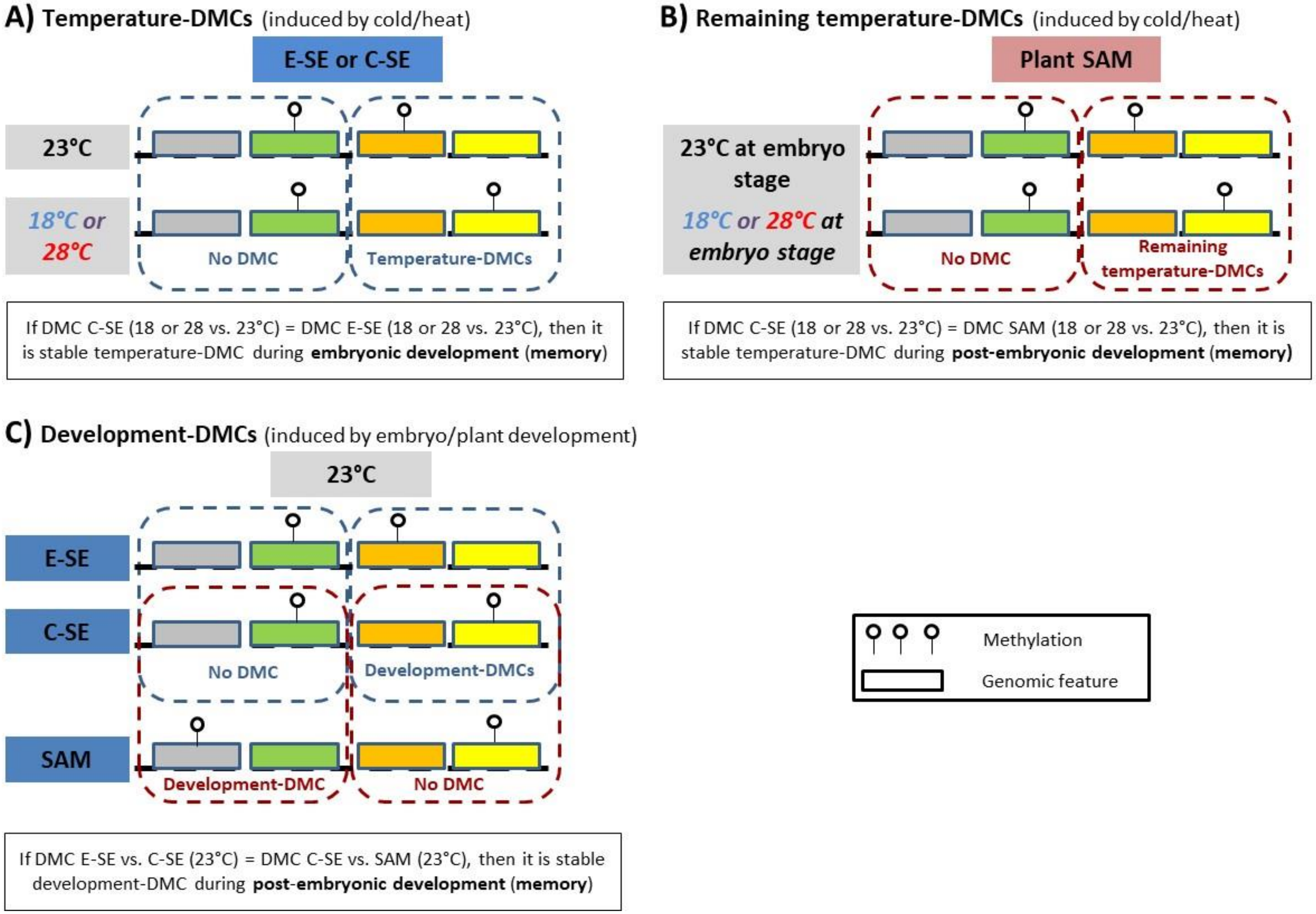


**Figure S6:** The different types of Differentially Methylated Cytosines **(**DMCs) and associated memory (stable, memorized DMCs during embryonic or post-embryonic development, see Fig. 5A).

**A.** DMCs detected in early (E-SE) or cotyledonary somatic embryos (C-SE) following temperature treatment (cold: 18 vs. 23°C; heat: 28 vs. 23°C) during the maturation phase. Stable temperature-DMCs from E-SE to C-SE contribute to temperature-induced embryonic memory.

**B.** DMCs detected in plant shoot apical meristem (SAM) following temperature treatment (cold: 18 vs. 23°C; heat: 28 vs. 23°C) during the embryo maturation phase. Stable, remaining temperature-DMCs from C-SE to SAM contribute to temperature-induced post-embryonic memory.

**C.** DMCs detected during embryonic (E-SE vs. C-SE) and post-embryonic (C-SE vs. SAM) development following embryo maturation at the reference temperature (23°C). Stable, remaining development-DMCs from embryonic to post-embryonic phase contribute to development-induced post-embryonic memory.

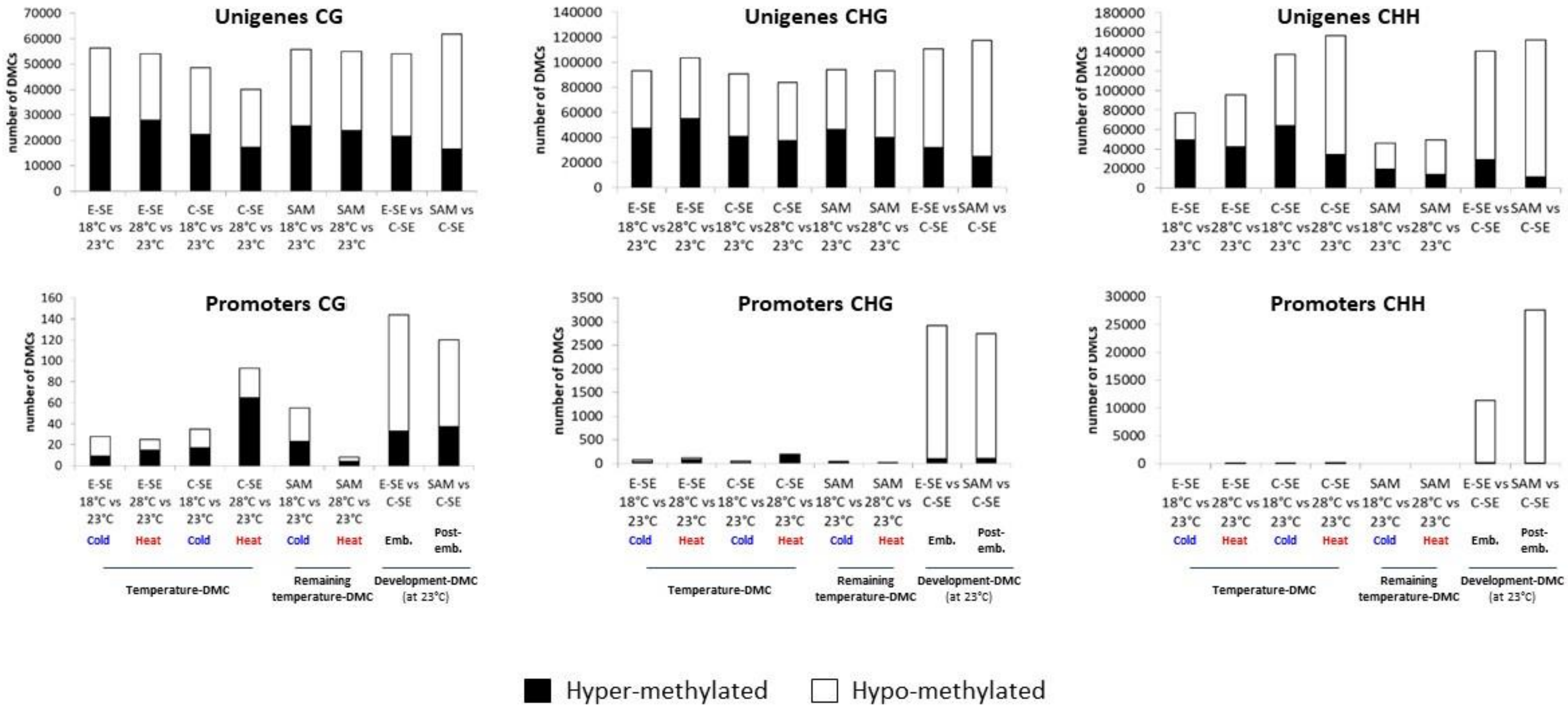


**Figure S7:** Number of hyper- and hypo-methylated DMCs for heat (28 vs. 23°C), cold (18 vs. 23°C), or embryonic (C-SE vs. E-SE) and post-embryonic (SAM vs. C-SE) developmental effects at 23°C in unigenes (up) and promoters (down) for the 3 methylation contexts (CG, CHG, CHH). The different DMC types (cold/heat temperature- and remaining temperature-DMCs as well as embryonic and post-embryonic development-DMCs) are indicated (see Fig. S6).

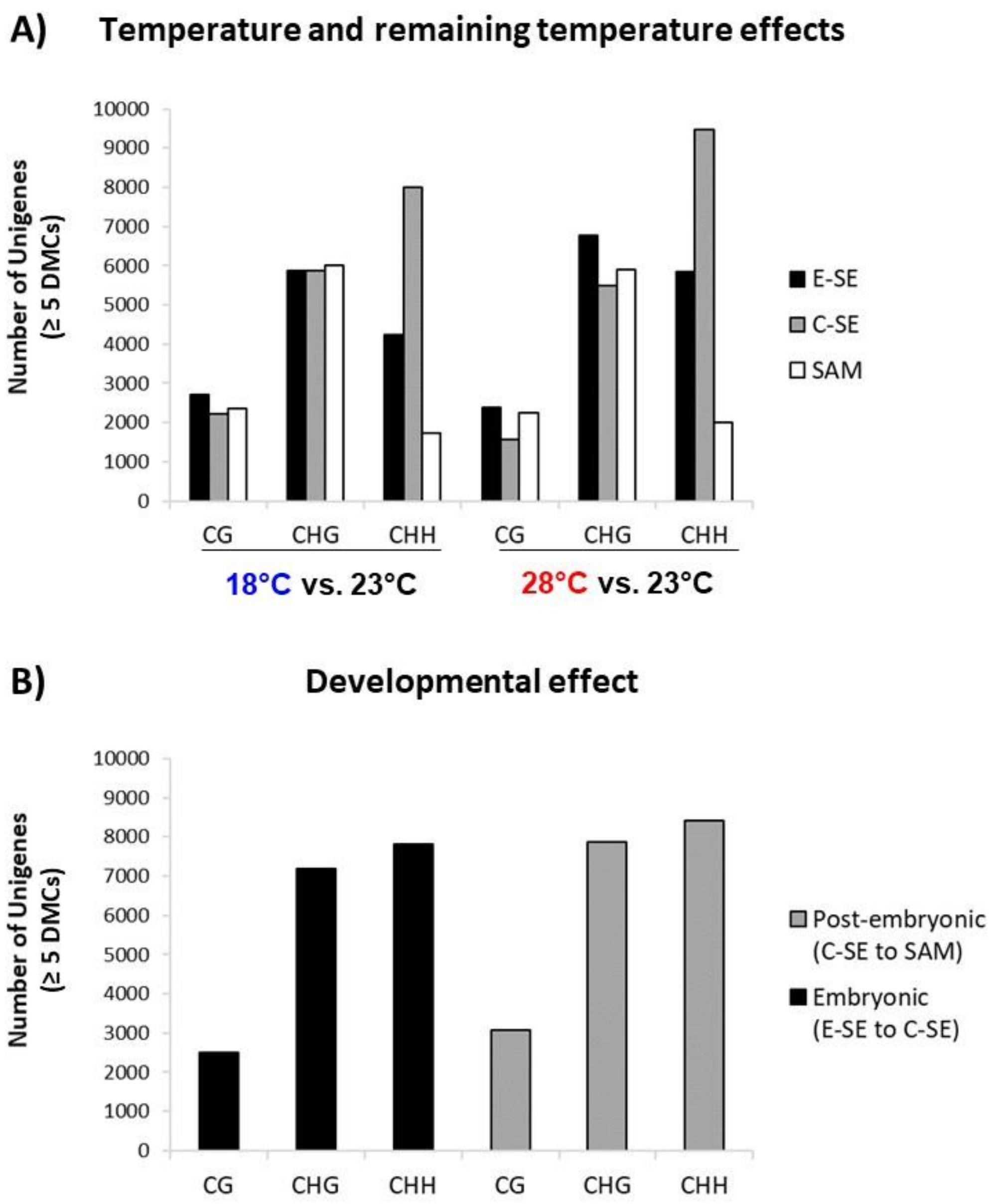


**Figure S8:** Number of unigenes with at least 5 DMCs for each context

**A.** Temperature effect (seen in E-SE, C-SE) and remaining temperature effect (seen in SAM) for cold (18 vs. 23°C) and heat (28 vs. 23°C).

**B.** Developmental effect (at 23°C) during embryonic (E-SE to C-SE) and post-embryonic development (C-SE to SAM).

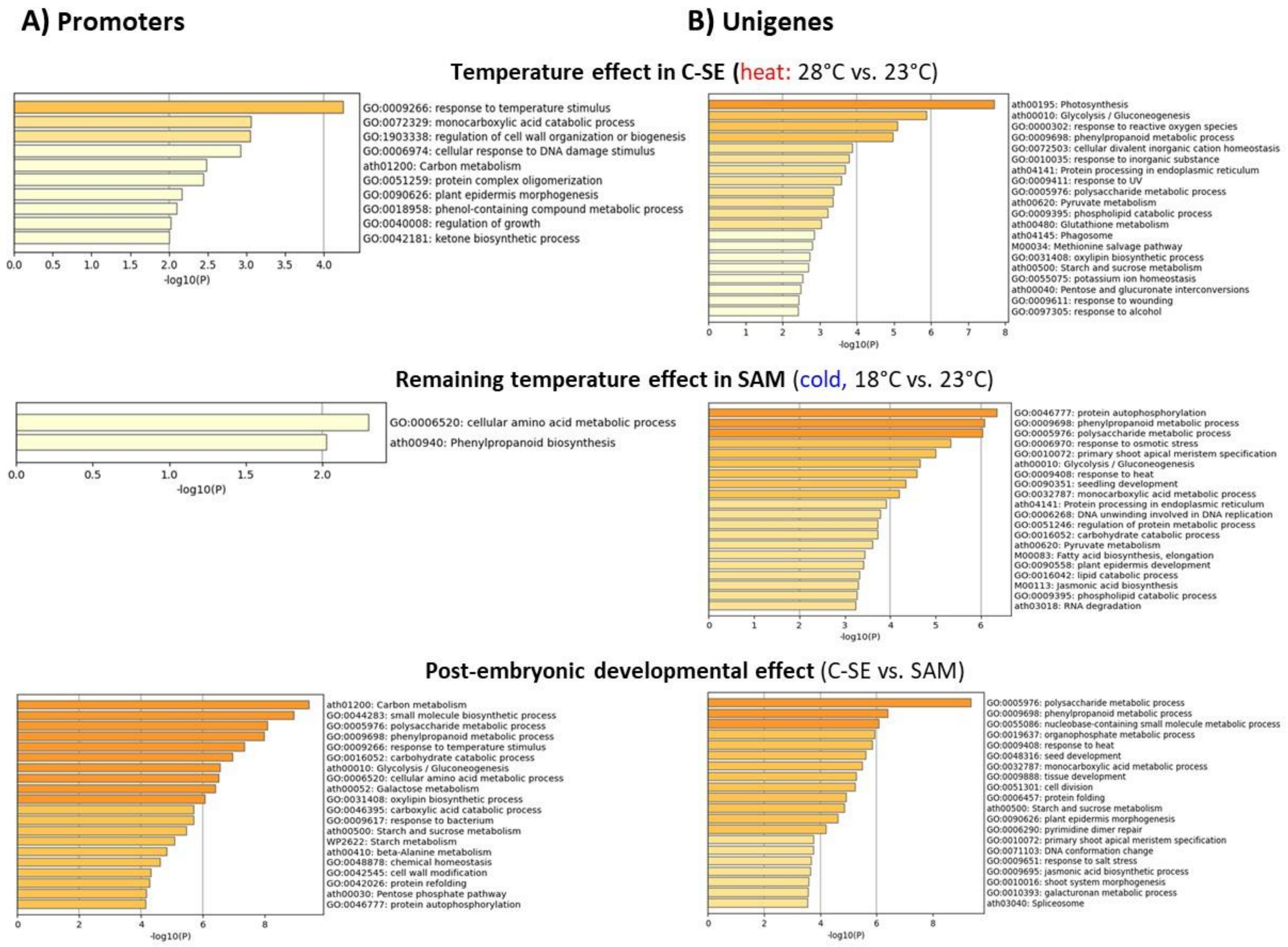


**Figure S9:** GO enrichment analysis based on promoters (**A**) and unigenes (**B**) with temperature-, remaining temperature- or development-DMCs considering all 3 methylation contexts (CG, CHG, CHH).

Dark orange bars highlight main molecular functions (-log10(P)>6). Only selected results are presented: heat effect in C-SE (28 vs. 23°C), cold effect in SAM (18 vs. 23°C) and post-embryonic developmental effect (at 23°C). See Mat. S3 for full GO enrichment analysis (including heat/cold effects in E-SE, cold effect in C-SE, heat effect in SAM and embryonic developmental effect).

**Supplementary Material**

**Material S1:** Phenotype characterization (biological and biochemical data).
**Material S2:** Designed probes for sequence capture bisulfite (SeqCapBis design, Roche).
**Material S3:** Gene Ontology (GO) enrichment analysis (Metascape).

Material S2 and S3 available on *data.gouv.fr*, the open platform for French public data (doi: 10.57745/PNGW7G).
